# Cortical control of virtual self-motion using task-specific subspaces

**DOI:** 10.1101/2019.12.13.862532

**Authors:** Karen E Schroeder, Sean M Perkins, Qi Wang, Mark M Churchland

## Abstract

Brain-machine interfaces (BMIs) for reaching have enjoyed continued performance improvements, yet there remains significant need for BMIs that control other movement classes. Recent scientific findings suggest that the intrinsic covariance structure of neural activity depends strongly on movement class, potentially necessitating different decode algorithms across classes. To address this, we developed a self-motion BMI based on cortical activity as monkeys performed non-reaching arm movements: cycling a hand-held pedal to progress along a virtual track. Unlike during reaching, we found no high-variance dimensions that directly correlated with to-be-decoded variables. Yet we could decode a single variable – self-motion – by non-linearly leveraging structure that spanned many high-variance neural dimensions. Resulting online BMI-control success rates approached those during manual control. These findings make two broad points regarding how to build decode algorithms that harmonize with the empirical structure of neural activity in motor cortex. First, even when decoding from the same cortical region (e.g., arm-related motor cortex) different movement classes may need to employ very different strategies. Although correlations between neural activity and hand velocity are prominent during reaching tasks, they are not a fundamental property of motor cortex and cannot be counted on to be present in general. Second, although one generally desires a low-dimensional readout, it can be beneficial to leverage a multi-dimensional high-variance subspace. Fully embracing this approach requires highly non-linear approaches tailored to the task at hand, but can produce near-native levels of performance.

**Significance Statement:** Many BMI decoders have been constructed for controlling movements normally performed with the arm. Yet it is unclear how these will function beyond the reach-like scenarios where they were developed. Existing decoders implicitly assume that neural covariance structure, and correlations with to-be-decoded kinematic variables, will be largely preserved across tasks. We find that the correlation between neural activity and hand kinematics, a feature typically exploited when decoding reach-like movements, is essentially absent during another task performed with the arm: cycling through a virtual environment. Nevertheless, the use of a different strategy, one focused on leveraging the highest-variance neural signals, supported high performance real-time BMI control.

## Introduction

Intracortical brain-machine interfaces (BMIs) that support reach-like tasks have proved successful in primates and human clinical trials (Ajiboye et al., 2017; Collinger et al., 2013; Ethier et al., 2012; Gilja et al., 2015, 2012; Shanechi et al., 2017; Shenoy and Carmena, 2014; Wodlinger et al., 2015). Yet it is unclear whether current decode algorithms will generalize well to non-reaching applications, even within the domain of arm movements. Early reach-based BMIs (Carmena et al., 2003; Chapin et al., 1999; Serruya et al., 2002; Taylor et al., 2002; Velliste et al., 2008; Wessberg et al., 2000) employed a strategy of inverting the ostensible neural encoding of kinematic variables, primarily hand velocity and direction. Despite evidence against literal kinematic encoding (Churchland et al., 2012; Michaels et al., 2016; Russo et al., 2018; Scott et al., 2001; Sergio et al., 2005; Sussillo et al., 2015), studies have continued to leverage robust correlations between neural activity and reach kinematics. Improvements have derived from honing this strategy (Gilja et al., 2012), from stabilizing the neural subspace containing the useful correlations (Degenhart et al., 2020; Gallego et al., 2020), and/or from better estimating the neural state (Aghagolzadeh and Truccolo, 2016; Makin et al., 2018). For example, Kao et al. (2015) leveraged dynamics, spanning many neural dimensions, to denoise activity in dimensions where correlations with reach kinematics were strong.

Thus, the vast majority of reach-related decode algorithms leverage the same statistical regularity: correlations between neural activity and hand velocity and/or position. The success of this approach is both motivated by (Chase and Schwartz, 2011; Shenoy et al., 2011) and serves to validate scientific assumptions. These include which external variables have robust correlations with neural activity (Taylor et al., 2002), whether activity evolves according to dynamics (Kao et al., 2015), and how neural responses adapt during learning (Ganguly et al., 2011; Sadtler et al., 2014; Taylor et al., 2002). A natural question is thus, can we still rely on the same scientific foundation when decoding movements of the arm that are not reach-like? Or is neural activity different enough that alternative strategies become necessary?

BMIs designed for different effectors, e.g. for tongue and speech decoding, have already had to contend with the possibility that strong correlations between neural activity and to-be-decoded variables may be absent, requiring highly non-linear approaches (Anumanchipalli et al., 2019; Liu et al., 2019). Even within the realm of reaching, performance is improved if a recurrent network precedes linear decoding of kinematics (Sussillo et al., 2016, 2012). This may be partly due to denoising (similar to Kao et al. 2015) but also suggests meaningful non-linear relationships. What then should we expect during non-reach-like arm movements? Will strong linear correlations with velocity persist, or will other more non-linear relationships become dominant? This question is intertwined with our evolving understanding of motor cortex covariance structure. During a given task, a handful of neural dimensions captures considerable variance (Churchland et al., 2007; Gallego et al., 2017; Sadtler et al., 2014). These high-variance dimensions are useful, because higher variance implies better noise resistance. Within a given task, neural covariance structure remains surprisingly fixed even when the decoder is altered (Golub et al., 2018; Sadtler et al., 2014) and can remain similar across related tasks (Gallego et al., 2018). If covariance remains fixed across very different tasks, then the nature of neural-vs-kinematic correlations would presumably be stable, as would the neural dynamics themselves.

Yet there is increasing experimental evidence that covariance is often not stable across tasks: covariance changes dramatically when cycling forward versus backward (Russo et al., 2018), when using one arm versus another (Ames and Churchland, 2019), when preparing versus moving (Elsayed et al., 2016; Inagaki et al., 2020; Kaufman et al., 2014), when reaching versus walking (Miri et al., 2017), and when co-contracting versus alternating muscle activity (Warriner et al., *under review*). Thus, in a new task there is no guarantee the high-variance dimensions will be the same or will show the same correlations with kinematics. Indeed, separating computations related to different tasks into different dimensions may be a common mechanism for neural populations to leverage different covariance patterns while preventing interference between them (Duncker et al., 2020). While this network strategy would impair the ability of fixed decoders to generalize across tasks, it could also allow the design of decoders that leverage task-specific relationships between neural activity and kinematic variables. To investigate, we employed a simple task in which monkeys cycle a hand-held pedal to move along a virtual track. The class of neural activity evoked by this task could potentially be leveraged by future BMI devices that guide self-motion. Furthermore, cycling is overtly different from reach- or joystick-based tasks. Prior explorations of self-motion-decoding borrowed from the strategies employed by reach-based BMIs, and decoded a whole-body directional vector (Rajangam et al., 2016) or classified the direction of a joystick intermediary (Libedinsky et al., 2016). This is a promising approach, but we wished to explore whether a task-specific approach might also be promising.

During cycling, activity that correlated with kinematics was relegated to low-variance dimensions. In contrast, there existed high-variance subspaces where neural activity had reliable non-linear relationships with intended self-motion. The ability to decode a one-dimensional command for virtual self-motion, from activity spanning many high-variance dimensions, produced both high accuracy and low latency. Success rates and acquisition times were close to those achieved under manual control. As a result, almost no training was needed; monkeys appeared to barely notice transitions from manual to BMI control. These findings make two points regarding how BMI decoding should interact with the basic properties of motor cortex activity. First, the neural relationships leveraged by traditional decoders are empirically reliable only during some behaviors. Second, even when those traditional relationships are absent, other task-specific relationships will be present and can support very accurate decoding.

## Methods

### Subjects and primary task

All procedures were approved by the Columbia University Institutional Animal Care and Use Committee. Subjects G and E were two adult male macaque monkeys (*Macaca mulatta*). Monkeys sat in a primate chair facing an LCD monitor (144 Hz refresh rate) that displayed a virtual environment generated by the Unity engine (Unity Technologies, San Francisco, CA). The head was restrained via a titanium surgical implant. While the monkey’s left arm was comfortably restrained, the right arm grasped a hand pedal. Cloth tape was used to ensure consistent placement of the hand on the pedal. The pedal connected via a shaft to a motor (Applied Motion Products, Watsonville, CA), which contained a rotary encoder that measured the position of the pedal with a precision of 1/10,000 of the cycle. The motor was also used to apply forces to the pedal, endowing it with virtual mass and viscosity.

Although our primary focus was on BMI performance, we also employed multiple sessions where the task was performed under manual control. Manual-control sessions allowed us to document the features of neural responses we used for decoding. They documented ‘normal’ performance, against which BMI performance could be compared. Monkey G performed eight manual-control sessions (average of 229 trials/session), each within the same day as one of the twenty BMI-controlled sessions (average of 654 trials/session). Monkey E performed five manual-control sessions (average of 231 trials/session) on separated days from the seventeen BMI-controlled sessions (average of 137 trials/session).

BMI-controlled sessions were preceded by 50 ‘decoder-training trials’ performed under manual control. These were used to train the decoder on that day. Decoder-training trials employed only a subset of all conditions subsequently performed during BMI control. Thus, BMI-controlled performance had to generalize to sequences of different distances not present during the decoder-training trials. Because they did not include all conditions, we do not analyze performance for decoder-training trials. All performance comparisons are made between manual-control and BMI-control sessions, which employed identical conditions, trial parameters, and definitions of success versus failure. We describe the task below from the perspective of manual control. The task was identical under BMI control except that position in the virtual environment was controlled by the output of a decoder rather than the pedal. We did not prevent or discourage the monkey from cycling during BMI-control, and he continued to do so as normal.

The cycling task required that the monkey cycle the pedal in the instructed direction to move through the virtual environment, and stop on top of lighted targets to collect juice reward. The color of the landscape indicated whether cycling must be ‘forward’ (green landscape, the hand moved away from the body at the top of the cycle) or ‘backward’ (tan landscape, the hand moved toward the body at the top of the cycle). In the primary task, cycling involved moving between stationary targets (in a subsequent section we describe an additional task used to evaluate speed control). There were 6 total conditions, defined by cycling direction (forward or backward) and target distance (2, 4, or 7 cycles). Distance conditions were randomized within same-direction blocks (3 trials of each distance per block), and directional blocks were randomized over the course of each session. Trials began with the monkey stationary on a target. A second target appeared in the distance. To obtain reward, the monkey had to cycle to that target, come to a halt ‘on top’ of it (in the first-person perspective of the task) and remain stationary for a hold period of 1000-1500 ms (randomized). A trial was aborted without reward if the monkey began moving before target onset (or in the 170 ms after, which would indicate attempted anticipation), if the monkey moved past the target without stopping, or if the monkey moved while awaiting reward. The next trial began 100 ms after the variable hold period. Monkeys performed until they received enough liquid reward that they chose to desist. As their motivation waned, they would at times take short breaks. For both manual-control and BMI-control sessions, we discarded any trials in which monkeys made no attempt to initiate the trial, and did not count them as ‘failed’. These trials occurred 2 ± 2 times per session (mean and standard deviation, Monkey G, maximum 10) and 3 ± 3 times per session (Monkey E, maximum 11).

For monkey G, an additional three manual-control sessions (189, 407, and 394 trials) were employed to record electromygraphic activity (EMG) from the muscles. We recorded from 5-7 muscles per session, yielding a total of 19 recordings. We made one or more recordings from the three heads of the *deltoid*, the lateral and long heads of *triceps brachii*, the *biceps brachii*, *trapezius,* and *latissimus dorsi*. These muscles were selected due to their clear activation during the cycling task.

### Surgery and neural/muscle recordings

Neural activity was recorded using chronic 96-channel Utah arrays (Blackrock Microsystems, Salt Lake City, UT), implanted in the left hemisphere using standard surgical techniques. In each monkey, an array was placed in the region of primary motor cortex (M1) corresponding to the upper arm. In monkey G, a second array was placed in dorsal premotor cortex (PMd), just anterior to the first array. Array locations were selected based on MRI scans and anatomical landmarks observed during surgery. Experiments were performed 1-8 months (monkey G) and 3-4 months (monkey E) after surgical implantation. Neural responses both during the task and during palpation confirmed that arrays were in the proximal-arm region of cortex.

Electrode voltages were filtered (band-pass 0.3 Hz – 7.5 kHz) and digitized at 30 kHz using Digital Headstages, Digital Hubs, and Cerebus Neural Signal Processors from Blackrock Microsystems. Digitized voltages were high-pass filtered (250 Hz) and spike events were detected based on threshold crossings. Thresholds were set to between −4.5 and −3 times the RMS voltage on each channel, depending on the array quality on a given day. On most channels, threshold crossings included clear action-potential waveforms from one or more neurons, but no attempt was made to sort action potentials.

Intra-muscular EMG recordings were made using pairs of hook-wire electrodes inserted with 30 mm x 27 gauge needles (Natus Neurology, Middleton, WI). Raw voltages were amplified and filtered (band-pass 10 Hz – 10 kHz) with ISO-DAM 8A modules (World Precision Instruments, Sarasota, FL), and digitized at 30 kHz with the Cerebus Neural Signal Processors. EMG was then digitally band-pass filtered (50 Hz – 5 kHz) prior to saving for offline analysis. Offline, EMG recordings were rectified, low-pass filtered by convolving with a Gaussian (standard deviation: 25 ms), downsampled to 1 kHz, and then fully normalized such that the maximum value achieved on each EMG channel was 1.

A real-time target computer (Speedgoat, Bern, CH) running Simulink Real-Time environment (MathWorks, Natick, MA) processed behavioral and neural data and controlled the decoder output in online experiments. It also streamed variables of interest to another computer that saved these variables for offline analysis. Stateflow charts were implemented in the Simulink model to control task state flow as well as the decoder state machine. Real-time control had millisecond precision.

Spike trains were causally converted to firing rates by convolving each spike with a beta kernel. The beta kernel was defined by temporally scaling a beta distribution (shape parameters: *α* = 3 and *β* = 5) to be defined over the interval [0, 275] ms and normalizing the kernel such that the firing rates would be in units of spikes/second. The same filtering was applied for online decoding and offline analyses. Firing rates were also mean centered (subtracting the mean rate across all times and conditions) and normalized. During online decoding, the mean and normalization factor were values that had been computed from the training data. We used soft normalization (Russo et al., 2018): the normalization factor was the firing rate range plus a constant (5 spikes/s).

### Computing trial-averaged firing rates

Analyses of BMI performance are based on real-time decoding during online performance, with no need to consider trial-averaged firing rates. However, we still wished to compute trial-averaged traces of neural activity and kinematics for two purposes. First, some aspects of decoder training benefited from analyzing trial-averaged firing rates. Second, we employ analyses that document basic features of single-neuron responses and of the population response (e.g., plotting neural activity within task-relevant dimensions). These analyses benefit from the denoising that comes from computing a time-varying firing rate across many trials. Due to the nature of the task, trials could be quite long (up to 20 cycles in the speed-tracking task), rendering the traditional approach of aligning all trials to movement onset insufficient for preserving alignment across all subsequent cycles. It was thus necessary to modestly adjust the time-base of each individual trial (e.g., stretching time slightly for a trial where cycling was faster than typical). We employed two alignment methods. Method A is a simplified procedure that was used prior to parameter fitting when training the decoder before online BMI control. This method aligns only times during the movement. Method B is a more sophisticated alignment procedure that was used for all offline analyses. This method aligns the entire trial, including pre- and post-movement data. For visualization, conditions with the same target distance (e.g., 7 cycles), but different directions, were also aligned to the same time base. Critically, any data processing that relied on temporal structure was completed in the original, unstretched time base prior to alignment.

Method A: The world position for each trial resembles a ramp between movement onset and offset. First, we identify the portion of each trial starting ¼ cycle into the movement and ending ¼ cycle before the end of the movement. We fit a line to the world position in this period and then extend that line until it intercepts the starting and ending positions. The data between these two intercepts is considered the movement data for each trial and is extracted. This movement data is then uniformly stretched in time to match the average trial length for each trial’s associated condition. This approach compresses slower than average movements and stretches faster than average movements within a condition, such that they can be averaged while still preserving many of the cycle-specific features of the data.

Method B: This method consists of a mild, non-uniform stretching of time in order to match each trial to a condition-specific template. For complete details, see Russo et al. 2018. (Russo et al., 2018)

### Neural variance captured analysis

Analysis of neural variance captured was based on successful trials from the three sessions, performed under manual control, with simultaneous neural and muscle recordings. Each session was split into a training set (250 trials for two of the sessions, 300 trials for the third) and testing set (100 trials for two of the sessions, 200 trials for the third). Neural dimensions were identified using a subset of training trials in order to match the size of the training sets used for online decoding (25 forward 7-cycle trials and 25 backward 7-cycle trials). Neural variance captured was computed (as described below) using the trial-averaged neural activity from the full training set for all conditions. We considered data from the full duration of each trial, including times before movement onset and after movement offset. We analyzed the variance captured by neural dimensions of three types. First, neural dimensions where activity correlated strongly with kinematic features. Second, neural dimensions where activity correlated strongly with muscle activity. Third, neural dimensions that captured robust ‘features’ leveraged by our decoder.

Dimensions of the third type were found as detailed in a dedicated section below. Dimensions of the first two types were found with ridge regression, using the model *z(r, t)* = *c* + *w^T^y(r, t)* where *z(r, t)* is the kinematic or muscle variable at time *t* during trial *r*, and *y(r, t)* is the corresponding *N*-dimensional vector of neural firing rates. The vector *w* defines a direction in neural space where activity correlates strongly with the variable *z*. It is found by minimizing the objective function 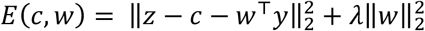, where *λ* is a free parameter that serves to regularize the weights *w*. We found multiple such vectors; e.g. *w_x−vel_* is a dimension where neural activity correlates with horizontal velocity and *w_biceps_* is a dimension where neural activity correlates with biceps activity. For each kinematic or muscle variable, we swept *λ* to find the value that yielded the largest coefficient of determination on held-out data, and then used that *λ* for all analysis. Because filtering of neural activity introduces a net lag, this analysis naturally assumes a ~100 ms lag between neural activity and the variables of interest. Results were extremely similar if we considered longer or shorter lags.

Before computing neural variance explained, each vector *w* was scaled to have unity norm, yielding a dimension in neural space. We wished to determine whether that dimension captured large response features that were reliable across trials. Thus, variance captured was always computed based on trial-averaged neural responses (averages taken across all trials in the data used to identify ***w***). We considered the matrix 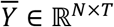 where *T* is the total number of time points across all conditions. Each row of 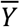 contains the trial-averaged firing rate of one neuron. We computed an *N* × *N* covariance matrix 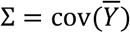 by treating rows of 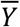 as random variables and columns as observations. The proportion of total neural variance captured by a given dimension, ***w***, is therefore:

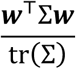

Some analyses considered the variance captured by a subspace spanned by a set of dimensions. To do so we took the sum of the variance captured by orthonormal dimensions spanning that space.

### Identifying neural dimensions

The response features leveraged by the decode algorithm are clearly visible in the top principal components of the population response, but individual features are not necessarily aligned with individual principal component dimensions. To find neural dimensions that cleanly isolated features, we employed dedicated preprocessing and dimensionality reduction approaches tailored to each feature. Dimensions were found based on the 50 decoder-training trials, collected at the beginning of each BMI-controlled session, before switching to BMI-control.

We sought a “moving-sensitive dimension” where activity reflected whether the monkey was stopped or moving. We computed spike-counts in time bins both when the monkey was moving (defined as angular velocity > 1 Hz) and when he was stopped (defined as angular velocity < .05 Hz). We ignored time bins that did not fall into either category. Spike counts were square-root transformed so that a Gaussian distribution becomes a more reasonable approximation (Thacker and Bromiley, 2001). This resulted, for each time-bin, in an activity vector (one element per recorded channel) of transformed spike counts and a label (stopped or moving). Linear discriminant analysis yielded a hyperplane that discriminated between stopped and moving. The moving-sensitive dimension, ***W**_move_*, was the vector normal to this hyperplane.

We sought four neural dimensions that captured rotational trajectories during steady-state cycling. Spike time-series were filtered to yield firing rates (as described above), then high-pass filtered (2^nd^ order Butterworth, cutoff frequency: 1 Hz) to remove any slow drift. Single-trial movement-period responses were aligned (Method A above) and averaged to generate an *N* × *T_f_* matrix 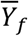 containing responses during forward cycling, and an *N* × *T_b_* matrix 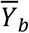 containing responses during backward cycling. We sought a 4-dimensional projection that would maximally capture rotational trajectories while segregating forward and backward data into different planes. Whereas the standard PCA cost function finds dimensions that maximize variance captured, we opted instead for an objective function that assesses the difference in variance captured during forward and backward cycling:

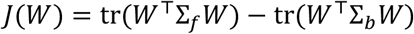

where 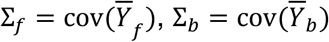, and *W* is constrained to be orthonormal. This objective function will be maximized when projecting the data onto *W* captures activity during forward cycling but not backward cycling. It will be minimized when the opposite is true: the projection captures activity during backward but not forward cycling. We thus define the forward rotational plane as the N × 2 matrix 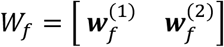 that maximizes *J*(*W*). Similarly, the backward rotational plane was the N x 2 matrix 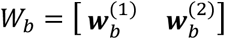 that minimizes *J*(*W*). Iterative optimization was used to find *W_f_* and *W_b_* using the Manopt toolbox (Boumal et al., 2014) as detailed in (Cunningham and Ghahramani, 2015). Optimization naturally results in *W_f_* and *W_b_* orthonormal to one another.

Finally, we sought dimensions where neural responses were maximally different, when comparing forward and backward cycling, in the moments just after movement onset. To identify the relevant epoch, we determined the time, *t_init_*, when the state machine would have entered the INIT state during online control. We then considered trial-averaged neural activity, for forward and backward cycling, from *t_init_* through *t_init_* + 200 ms. We applied PCA and retained the top three dimensions: 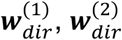, and 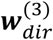. Such dimensions capture both how activity evolves during the analyzed epoch, and how it differs between forward and backward cycling.

### Computing probability of moving

Based on neural activity in the moving-sensitive dimension, an HMM inferred the probability of being in each of two behavioral states: ‘moving’ or ‘stopped’ (Kao et al., 2017a). As described above, square-rooted spike counts in the decoder-training data were already separated into ‘moving’ and ‘stopped’ sets for the purposes of identifying ***W**_move_*. We projected those counts onto ***W**_move_* and a fit a Gaussian distribution to each set.

When under BMI control, we computed *p_move_*(*t*), the probability of being in the ‘moving’ state, given the entire sequence of current and previously observed square-rooted spike counts. *p_move_*(*t*) was computed with a recursive algorithm that used the state transition matrix

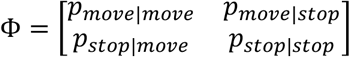

and knowledge of the Gaussian distributions. Φ encodes assumptions about the probability of transitioning from one state to the next at any given bin. For each monkey, we chose reasonable values for Φ based on preliminary data. For monkey G, we set *p_move|stop_* = .0001 and *p_stop|move_* = .002. For monkey E, we set *p_move|stop_* = .0002 and *p_stop|move_* = .004. These values were used for all BMI-controlled sessions.

### Computing steady-state direction and speed

During BMI control, we wished to infer the neural state in the four ‘rotational’ dimensions, spanned by [*W_f_*, *W_b_*]. We began with *y_t_*, a vector containing the firing rate of every neuron at the present time *t*, computed by causally filtering spikes and preprocessing as described above. For this computation only, computation of *y_t_* included high-pass filtering to remove any drift on timescales slower than the rotations (same filter used when identifying *W_f_* and *W_b_*). We applied a Kalman filter of the form

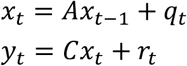

where 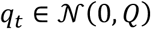, and 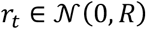. In these equations, *x_t_* represents the true underlying neural state in the rotational dimensions and the firing rates *y_t_* are a noisy reflection of that underlying state. We employed filtered firing rates, rather than binned spike-counts, for purely opportunistic reasons: it consistently yielded better performance.

The parameters of the Kalman filter were fit based on data from the decoder-training trials. We make the simplifying assumption that the ‘ground truth’ state is well-described by the trial-averaged firing rates projected onto [*W_f_*, *W_b_*]. If so, *A* and *Q* can be directly inferred from the evolution of that state, and *R* reflects to the degree to which single-trial firing rates differ from their idealized values:

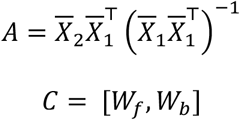

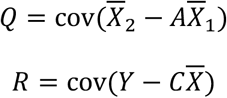

where

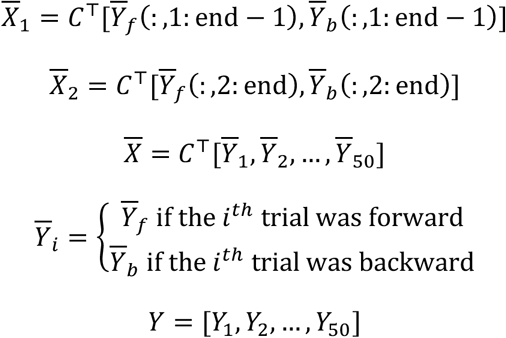

where 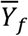 and 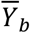 are the trial-averaged activity when cycling forward and backward respectively, *Y_i_* is the neural activity for the *i*-th trial in the training set, and indexing uses Matlab notation. Online inference yields an estimate 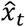 at each millisecond *t*, computed recursively using the steady-state form of the Kalman filter (Malik et al., 2011).

Angular momentum was computed in each plane as the cross product between the estimated neural state and its derivative, which (up to a constant scaling) can be written

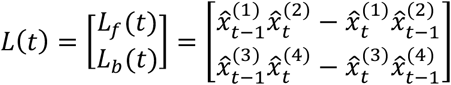

where the superscript indexes the elements of 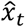. We fit two-dimensional Gaussian distributions to these angular momentums for each of three behaviors in the training data: ‘stopped’ (speed < .05 Hz), ‘pedaling forward’ (velocity > 1 Hz), and ‘pedaling backward’ (velocity < −1 Hz). Online, the likelihood of the observed angular momentums with respect to each of these three distributions dictated the steady-state estimates of direction and speed. We’ll denote these three likelihoods *f_stop_*, *f_forward_*, and *f_backward_*.

In principle, one could render a simple three-valued decode (stopped, moving forward, moving backward) based on the highest likelihood. However, we wanted the decoder to err on the side of withholding movement, and for movement speed to reflect certainty regarding direction. We set *speed_steady_* to zero unless the value of *f_stop_* was below a conservative threshold, set to correspond to a Mahalanobis distance of 3 between *L* and the distribution of angular momentums when stopped. (In one dimension this would be equivalent to being more than three standard deviations from the mean neural angular momentum when stopped). When *f_stop_* was below threshold, we decoded direction and speed as follows:

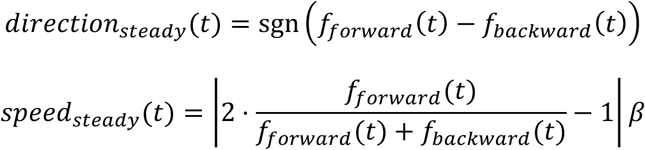

where 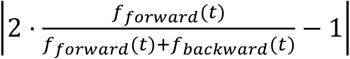 varies between 0 and 1 depending on the relative sizes of the likelihoods (yielding a slower velocity if the direction decode is uncertain) and *β* is a direction-specific constant whose purpose is simply to scale up the result to match typical steady-state cycling speed. In practice, certainty regarding direction was high at most moments and *speed_steady_* was thus close to the maximal value set by *β*.

### Computing initial direction and speed

Decoded motion was determined by a state-machine with four states: STOP, INIT, EARLY, and STEADY. Transitions between these states were determined primarily by *p_move_*. Decoded velocity was zero for STOP and INIT and was determined by *speed_steady_* (as described above) when in STEADY. Because rotational features were not yet robustly present in the EARLY state, we employed a different method to infer initial direction and speed. Both were determined at *t_early_*, the moment the EARLY state was entered. These values then persisted throughout the remainder of the EARLY state. Decoded direction was determined by projecting the vector of firing rates, at the moment the EARLY state was entered, onto the three initial-direction dimensions, 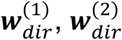, and 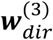. These directions were computed based on decoder-training trials (see above) as were the Gaussian distributions of single-trial projections when cycling forward versus backward (using the moment the EARLY state would have been entered). This allowed us to compute the likelihoods, *g_forward_* and *g_backward_*, of the present projection given each distribution. If the observed projection was not an outlier (>10 Mahalanobis distance units) with respect to both distributions, initial direction and speed were computed as:

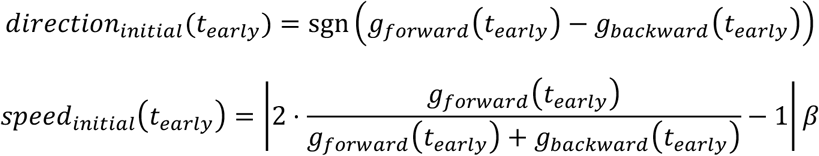

If the observed neural state was an outlier, we assumed rotational structure was likely already present and initial direction and speed were set to *direction_steady_* and *speed_steady_* as described above.

### Smoothing of decoded velocity

The decoder state machine produced an estimate of velocity, *v_dec_*, at every millisecond. During the STOP and INIT states, this estimate was always zero and the monkey’s position in the virtual environment was held constant. During the EARLY and STEADY states, this estimate was smoothed with a trailing average:

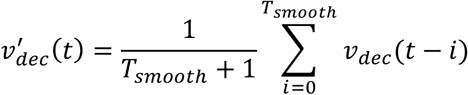

where *T_smooth_* = min(500, *t* − *t_early_*), i.e., the trailing average extended in history up to 500 ms or to the moment the EARLY state was entered, whichever was shorter. 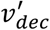 was integrated every millisecond to yield decoded position in the virtual environment. In the speed-tracking experiment (described below) there was no need to smooth *v_dec_* prior to integration because the speed estimate had already been smoothed.

### Speed-tracking task

In addition to the primary task (where the monkey traveled 2-7 cycles between stationary targets) we employed a speed-tracking task, in which the monkey was required to match his virtual speed to an instructed speed. Speed was instructed implicitly, via the relative position of two moving targets. The primary target was located a fixed distance in front of the monkey’s location in virtual space: the secondary target fell ‘behind’ the first target when cycling was too slow, and pulled ‘ahead’ if cycling was too fast. This separation saturated for large errors, but for small errors was proportional to the difference between the actual and instructed speed. This provided sufficient feedback to allow the monkey to track the instructed speed even when it was changing. Because there was no explicit cue regarding the absolute instructed speed, monkeys began cycling on each trial unaware of the true instructed speed profile and ‘honed in’ on that speed over the first ~2 cycles.

We quantify instructed speed not in terms of the speed of translation through the virtual environment (which has arbitrary units) but in terms of the physical cycling velocity necessary to achieve the desired virtual speed. E.g., an instructed speed of 2 Hz necessitated cycling at an angular velocity of 2 Hz to ensure maximal reward. Under BMI control, the output of the decoder had corresponding units. For example, a 2 Hz angular velocity of the neural trajectory produced movement at the same speed as 2 Hz physical cycling (see *‘Neural features for speed-tracking’* for details of decoder). Reward was given throughout the trial so long as the monkey’s speed was within 0.2 Hz of the instructed speed. We employed both constant and ramping instructed-speed profiles.

Constant profiles were at either 1 Hz or 2 Hz. Trials lasted 20 cycles. After 18 cycles, the primary and secondary targets (described above) disappeared and were replaced by a final stationary target two cycles in front of the current position. Speed was not instructed during these last two cycles; the monkey simply had to continue cycling and stop on the final target to receive a large reward. Analyses of performance were based on the ~16 cycle period starting when the monkey first honed in on the correct speed (within 0.2 Hz of the instructed speed) and ending when the speed-instructing cues disappeared 2 cycles before the trial’s end.

Ramping profiles began with three seconds of constant instructed speed to allow the monkey to hone in on the correct initial speed. Instructed speed then ramped, over 8 seconds, to a new value, and remained constant thereafter. As for constant profiles, speed-instructing cues disappeared after 18 cycles and the monkey cycled two further cycles before stopping on a final target. Again, analyses of performance were based on the period from when the monkey first honed in on the correct speed, to when the speed-instructing cues disappeared. There were two ramping profiles: one ramping up from 1 to 2 Hz, and one ramping down from 2 to 1 Hz. There were thus four total speed profiles (two constant and two ramping). These were performed for both cycling directions (presented in blocks and instructed by color as in the primary task) yielding eight total conditions. This task was performed by monkey G, who completed an average of 166 trials/session over 2 manual-control sessions and an average of 116 trials/session over 3 BMI-control sessions. BMI-control sessions were preceded by 61-74 ‘decoder-training trials’ performed under manual control. Decoder-training trials employed the two constant speeds and not the ramping profiles. Thus, subsequent BMI-controlled performance had to generalize to these situations.

As will be described below, the speed decoded during BMI control was low-pass filtered to remove fluctuations due to noise. This had the potential to actually make the task easier under BMI control, given that changes in instructed speed were slow within a trial (excepting the onset and offset of movement). We did not wish to provide BMI control with an ‘unfair’ advantage in comparisons with manual control. We therefore also low-pass filtered virtual speed while under manual control. Filtering (exponential, τ = 1 second) was applied only when speed was above 0.2 Hz, so that movement onset and offset could remain brisk. This aided the monkey’s efforts to track slowly changing speeds under manual control.

In manual-control sessions, trials were aborted if there was a large discrepancy between actual and instructed speed. This ensured that monkeys tried their best to consistently match speed at all times. A potential concern is that this could also mask, under BMI control, errors that would have been observed had the trial not aborted. To ensure that such errors were exposed, speed discrepancies did not cause trials to abort when under BMI control. This potentially puts BMI performance at a disadvantage relative to manual control, where large errors could not persist. In practice this was not an issue as large errors were rare.

### Neural features for speed-tracking

Although the speed-tracking experiment leveraged the same dominant neural responses that were used in the primary experiment, some quantities were computed slightly differently. These changes reflected a combination of small improvements (the speed-tracking task was performed after the primary experiment described above) and modifications to allow precise control of speed. The probability of moving, *p_move_*, was calculated using a different set of parameters, largely due to changes in recording quality in the intervening time between data collection from the primary experiment and data collection for the speed-tracking experiment. The bin size was increased to 100 ms and the following state transition values were used: *p_move|stop_* = .0005 and *p_stop|move_* = .0005. In addition, we observed that the square-root transform seemed to be having a negligible impact at this bin size, so we removed it for simplicity. To avoid losing rotational features at slower speeds, we dropped the cutoff frequency of the high-pass filter, applied to the neural firing rates, from 1 Hz to 0.75 Hz.

In computing *direction_steady_*, the same computations were performed as for the primary-experiment, with one exception: a new direction was not necessarily decoded every millisecond. In order to decode a new direction, the follow conditions needed to be met: 1) the observed angular momentums had a Mahalanobis distance of less than 4 to the distribution corresponding to the decoded direction, 2) the observed angular momentums had a Mahalanobis distance of greater than 6 to the distribution corresponding to the opposite direction. These criteria ensured that a new steady-state direction was only decoded when the angular momentums were highly consistent with a particular direction. When these criteria were not met, the decoder continued to decode the same direction from the previous time step. This improved decoding but did not prevent the decoder from accurately reversing direction when the monkey stopped and reversed his physical direction (e.g., between a forward and backward trial).

Speed was computed identically in the EARLY and STEADY states, by decoding directly from the rotational plane corresponding to the decoded direction. A coarse estimate of speed was calculated as the derivative of the phase of rotation:

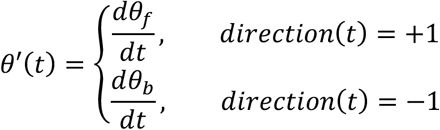

where *θ_f_ (t)* and *θ_b_ (t)* are the phases within the two planes of the neural state estimate 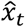, *direction* corresponds to *direction_early_* while in the EARLY state and *direction_steady_* while in the STEADY state, and the derivative *θ′* is computed in units of Hz. The coarse speed estimate, *θ*’, was then smoothed with an exponential moving average (τ = 500 ms) to generate the variable *speed*. Saturation limits were set such that, when moving, *speed* never dropped below 0.5 Hz or exceeded 3.5 Hz, to remain in the range at which monkeys can cycle smoothly. On entry into EARLY or STEADY from either INIT or EXIT, *speed* was reset to a value of 1.5 Hz. Thus, when starting to move, the initial instantaneous value of *speed* was set to an intermediate value and then relaxed to a value reflecting neural angular velocity.

The speed-tracking task employed two new conditions for decoder state transitions. First, transitions from INIT to EARLY required that a condition termed “confident initial direction decode” was obtained. This condition was met when the Mahalanobis distance from the neural state in the initial-direction subspace to either the forward or backward distributions dropped below 4. Second, transitions into the EXIT state required (in addition to a drop in *p_move_*) that the observed angular momentums, *L*, belong to a set termed ‘Stationary’. This set was defined as all *L* with a Mahalanobis distance of less than 4 to the ‘stopped’ distribution of angular momentums, which was learned from the training set.

## Results

### Behavior

We trained two monkeys (G and E) to rotate a hand-held pedal to move through a virtual environment (**Fig. 1**). All motion was along a linear track – no steering was necessary. Consistent with this, a single pedal was cycled with the right arm only. Our goal when decoding was to reconstruct the virtual self-motion produced by that single pedal. On each trial, a target appeared in the distance. To acquire that target, monkeys produced virtual velocity in proportion to the rotational velocity of the pedal. The color of the environment (lush and green versus desert-like and tan) instructed cycling direction. When the environment was green (**Fig. 1a**, *left*) forward virtual motion was produced by cycling ‘forward’ (i.e., with the hand moving away from the body at the top of the cycle). When the environment was tan (**Fig. 1a**, *right*) forward virtual motion was produced by cycling ‘backward’ (the hand moving toward the body at the top of the cycle). Cycling in the wrong direction produced motion away from the target. Trials were presented in blocks of forward or backward trials. Within each block, targets were separated by a randomized distance of 2, 4 or 7 cycles. Acquisition of a target was achieved by stopping and remaining stationary ‘on top’ of the virtual target for a specified time. Reward was then given and the next target appeared.

**Figure 1.**
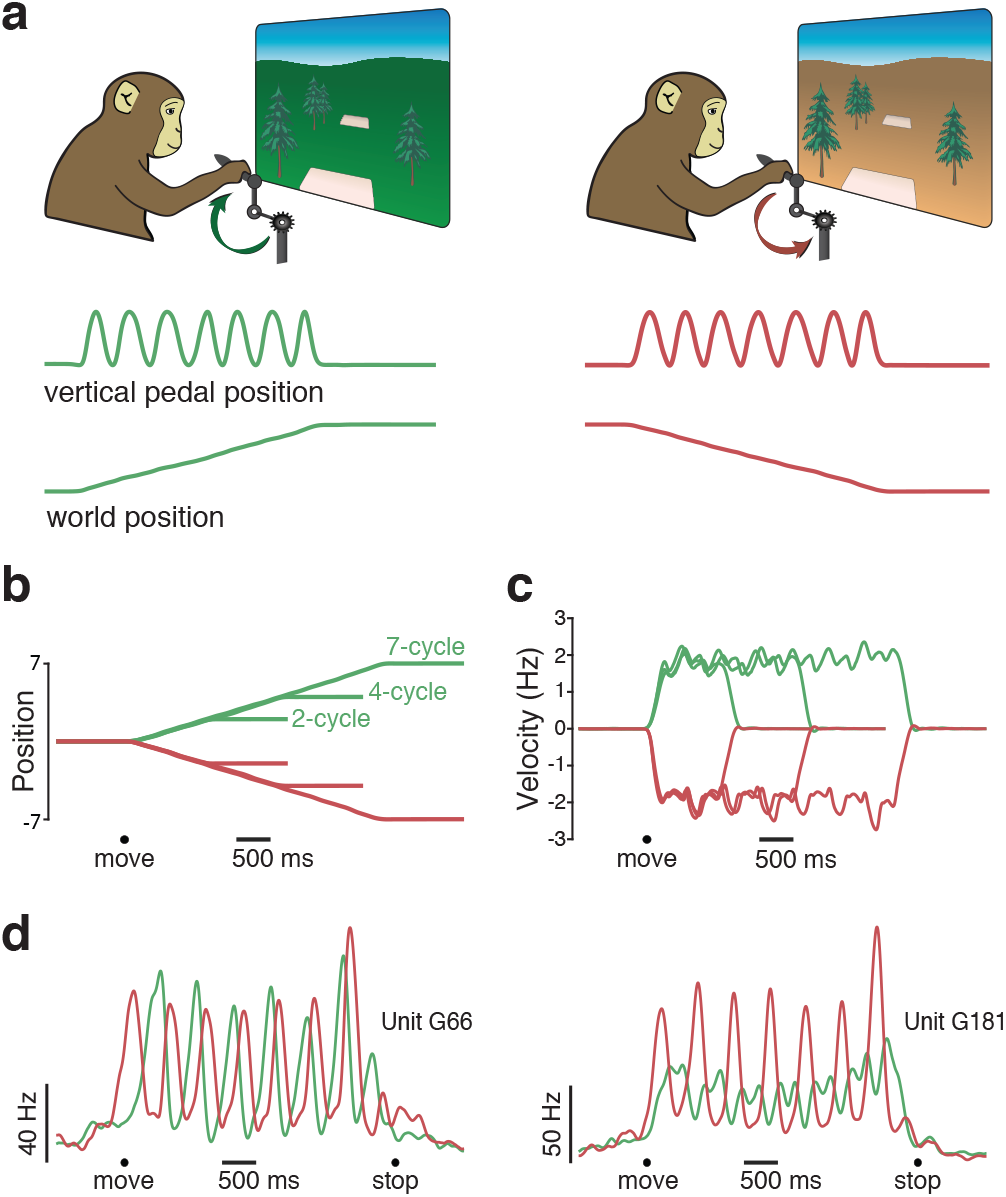
A cycling task that elicits rhythmic movements. **(a)** Monkeys rotated a hand-held pedal forward (*left*, cued by a green background) or backward (*right*, cued by a tan background) to progress through a virtual environment. Traces at bottom plot pedal kinematics (vertical position) and the resulting virtual world position for two example manual-control trials. On both of these trials (one forward and one backward) the monkey progressed from one target to another by cycling seven cycles. **(b)** Trial-averaged virtual position from a typical manual-control session. Each trace plots the change in virtual position (from a starting position of zero) for one of six conditions: forward or backward for 2, 4, or 7 cycles. Black circle indicates the time of movement onset. Trials were averaged after being aligned to movement onset, and then scaled such that the duration of each trial matched the average duration for that condition. **(c)** Trial-averaged pedal rotational velocity from the same session, for the same six conditions. **(d)** Firing rates of two example units. Trial-averaged firing rates (computed after temporally aligning trials) are shown for two conditions: forward (*green*) and backward (*red*) for seven cycles. Black circles indicate the timing of movement onset and offset.

Monkeys performed the task well, moving swiftly between targets, stopping accurately on each target, and remaining stationary until the next target was shown. Monkeys cycled at a pace that yielded nearly linear progress through the virtual environment (**Fig. 1b**). Although not instructed to cycle at any particular angular velocity, monkeys adopted a brisk ~2 Hz rhythm (**Fig. 1c**). Small ripples in angular velocity were present during steady-state cycling; when cycling with one hand it is natural for velocity to increase on the downstroke and decrease on the upstroke. Success rates were high, exceeding 95% in every session (failures typically involved over- or under-shooting the target location). This excellent performance under manual control provides a stringent bar by which to judge performance under BMI control.

BMI control was introduced after monkeys were adept at performing the task under manual control. Task structure and the parameters for success were unchanged under BMI control, and no cue was given regarding the change from manual to BMI control. For a BMI session, the switch to BMI control was made after completion of 50 decoder-training trials performed with manual-control (25 forward and 25 backward 7-cycle trials). The decoder was trained on these trials, at which point the switch was made to BMI control for the remainder of the session. For monkey G, we occasionally included a session of manual-control trials later in the day to allow comparison between BMI and manual performance. For Monkey E we used separate (interleaved) sessions to assess manual-control performance, because he was willing to perform fewer total trials per day.

During both BMI control and manual control, the monkey’s ipsilateral (non-cycling) arm was restrained. The contralateral (cycling) arm was never restrained. We intentionally did not dissuade the monkey from continuing to physically cycle during BMI control. Indeed, our goal was that the transition to BMI control would be sufficiently seamless to be unnoticed by the monkey, such that he would still believe that he was in manual control. An advantage of this strategy is that we are decoding neural activity when the subject attempts to actually move, as a patient presumably would. Had we insisted the arm remain stationary, monkeys would have needed to actively avoid patterns of neural activity that drive movement – something a patient would not have to do. Allowing the monkey to continue to move normally also allowed us to quantify decoder performance via direct comparisons with intended (i.e., actual) movement. This is often not possible when using other designs. For example, in Rajangam et. al. (Rajangam et al., 2016), performance could only be assessed via indirect measures (e.g., time to target) because what the monkey was actually intending to do at each moment was unclear. We considered these advantages to outweigh a potential concern: a decoder could potentially ‘cheat’ by primarily leveraging activity driven by proprioceptive feedback (which would not be present in a paralyzed patient). This is unlikely to be a large concern. Recordings were made from motor cortex, where robust neural responses precede movement onset. Furthermore, we have documented that motor cortex population activity during cycling is quite different from that within the proprioceptive region of primary somatosensory cortex (Russo et al., 2018). Thus, while proprioceptive activity is certainly present in motor cortex (Fetz et al., 1980; Lemon et al., 1976; Schroeder et al., 2017; Suminski et al., 2009) especially during perturbations (Pruszynski et al., 2011), the dominant features of M1 activity that we leverage are unlikely to be primarily proprioceptive.

Our goal was to use healthy animals to determine strategies for leveraging the dominant structure of neural activity. This follows the successful strategy of BMI studies that leveraged the well-characterized structure of activity during reaching. Of course, the nature of the training data used to specify decode parameters (e.g., the weights defining the key neural dimensions) will necessarily be different for a healthy animal that cannot understand verbal instructions and an impaired human that can. We thus stress that our goal is to determine a robust and successful decode strategy that works in real time during closed-loop performance. We do not attempt to determine the best approach to parameter specification, which in a patient would necessarily involve intended or imagined movement.

### Neural activity and decoding strategy

We recorded motor cortical activity using 96-channel Utah arrays. For monkey G, one array was implanted in primary motor cortex (M1) and a second in dorsal premotor cortex (PMd). For monkey E, a single array was implanted in M1. For each channel we recorded times when the voltage crossed a threshold. Threshold crossings typically reflected individual spikes from a small handful of neurons (a multi-unit). Spikes from individual neurons could be clearly seen on many channels, but no attempt was made to spike-sort. The benefit of sorting is typically modest when controlling a prosthetic device (Christie et al., 2014), and reduced-dimension projections of motor cortex population activity are similar whether based on single or multi-units (Trautmann et al., 2019). Unit activity was strongly modulated during cycling (**Fig. 1d**). The phase, magnitude, and temporal pattern of activity depended on whether cycling was stopped, moving forward (*green* traces) or moving backward (*red* traces). A key question is how these unit-level features translate into population-level features that might be leveraged to estimate intended motion through the virtual environment.

In traditional decoding approaches (**Fig. 2a**) neural activity is hypothesized (usefully if not literally) to encode kinematic signals, which can be decoded by inverting the encoding scheme. Although nonlinear methods (e.g., Kalman filtering of the neural state) are often used to estimate neural activity, the final conversion to a kinematic command is typically linear or roughly so. To explore kinematic encoding in the present task, we used ridge regression to identify neural dimensions in which activity correlated well with kinematics on single trials. For each kinematic variable, the degree of regularization was chosen to maximize generalization R^2^ – how well the kinematic variable could be reconstructed from activity in that neural dimension (**Fig. 2b;** *orange symbols*). We then computed the neural variance captured – the magnitude of the neural signals in that dimension (**Fig. 2b;** *dark teal bars*). The dimensions that best reconstructed kinematic signals all captured very little neural variance (**Fig. 2b**). The neural variance captured could not be increased via regularization without decreasing generalization R^2^ (i.e., without reducing the correlation with kinematics). These findings were also true for dimensions found to correlate well with muscle activity (**Fig. 2b**, *light teal bars).*

**Figure 2.**
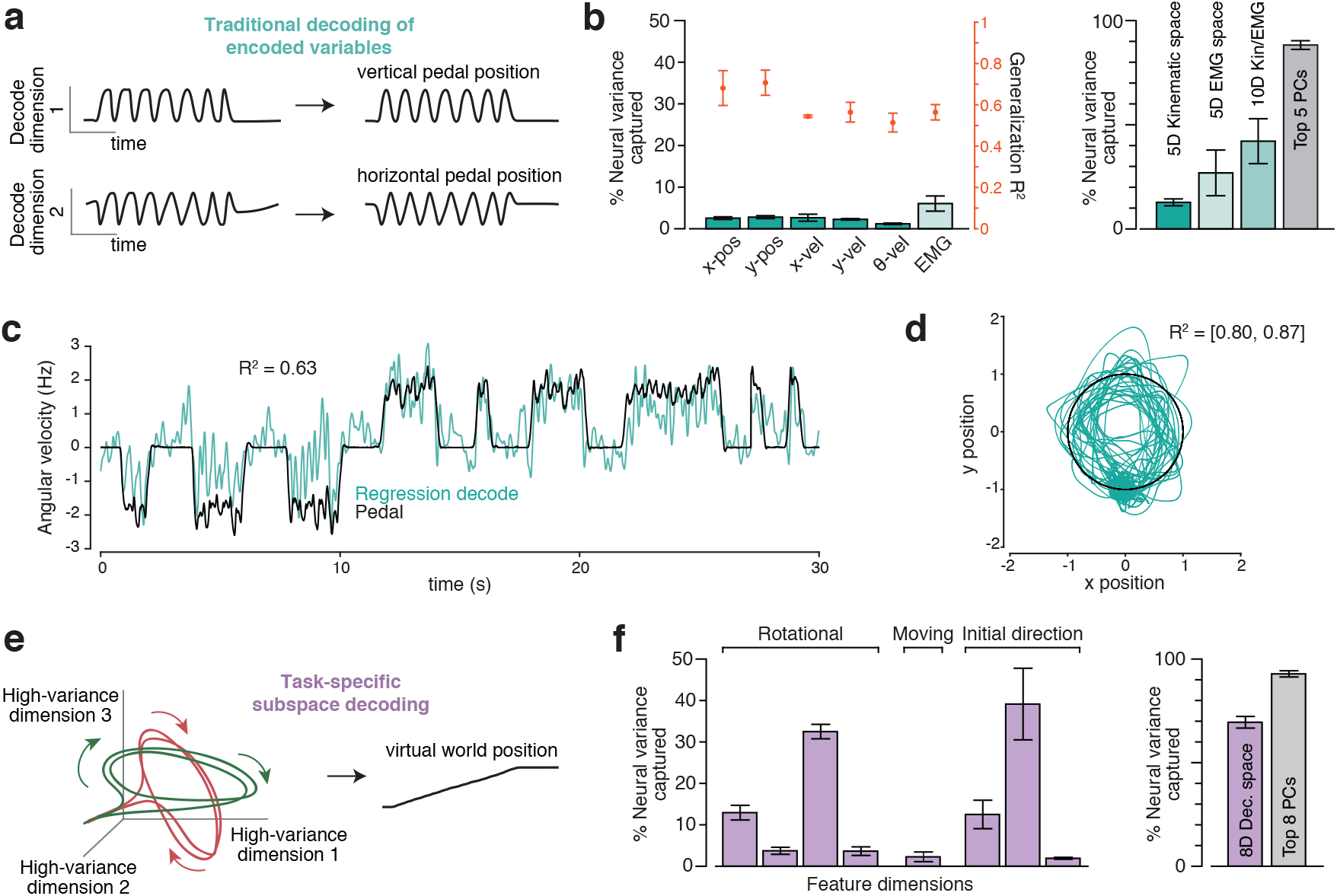
Different decode strategies leverage neural signals with different magnitudes. **(a)** In the traditional decoding strategy, neural firing rates are assumed to predominantly encode the key variables. The encoding model is usually assumed to be roughly linear when variables are expressed appropriately. For example, cosine tuning for reach velocity is equivalent to a linear dependence on horizontal and vertical velocity. The goal of decoding is to invert encoding. Thus, decoding dimensions should capture the dominant signals in the neural data (because those are what is encoded). **(b)** Variance of the neural population response captured by dimensions used to decode kinematic parameters (*teal bars*) and muscle activity (*light teal bar*). *Left subpanel*: neural variance captured (left axis) for dimensions correlating with kinematic variables (individual variables shown separately) and muscles (average across 19 recordings, standard error computed across recordings). Also shown are the associated generalized R2 values (right axis) for each decoder. *Right subpanel*: total variance captured by subspaces spanned by kinematic-decoding dimensions, muscle-decoding dimensions, or both. (These are not the sum of the individual variances as dimensions were not always orthogonal). We had different numbers of EMG recordings per day and thus always selected a subset of five. Variance captured by the top five principal components is shown for comparison. In both subpanels, data are from three manual-control sessions where units (192 channels per day) and muscles (5-7 channels per day) were recorded simultaneously. Each bar/point with error bars plots the average and standard error across sessions. **(c)** Example cross-validated regression performance for offline decoding of angular velocity. R^2^ is the coefficient of determination for the segment of data shown. **(d)** Example cross-validated regression performance for offline decoding of horizontal and vertical pedal position. R^2^ is the coefficient of determination for the segment of data shown, same time period as (c). **(e)** A new strategy that can be applied even if the dominant signals do not have the goal of encoding. This strategy seeks to find neural response features that have a robust relationship with the variable one wishes to decode. That relationship may be complex or even incidental, but is useful if it involves high-variance response features. **(f)** Similar plot to (b) but for the dimensions upon which our decoder was built. *Left subpanel*: neural variance captured for each of these eight dimensions. *Right subpanel*: neural variance captured by the eight-dimensional subspace spanned by those dimensions. Variance captured by the top eight principal components is shown for comparison.

The strikingly low (less than 5%) neural variance captured in kinematic-correlating dimensions was initially surprising, since single-neuron responses were robustly sinusoidally modulated, just like many kinematic variables. Yet sinusoidal response features were often superimposed upon other response features (e.g., overall shifts in rate when moving versus not moving). Sinusoidal features also displayed phase relationships, across forward and backward cycling, that were inconsistent with kinematic encoding. As a result, dimensions where activity correlated strongly with kinematics captured little response variance. This underscores that, when correlations are incidental, one cannot count on them being consistently present across behaviors. Furthermore, which correlations are more prevalent can also change. During reaching, correlations are typically strongest with velocity (a consequence of phasic activity) while in the present case correlations with position were at least as strong as those with velocity, both in terms of generalization R^2^ and in terms of neural variance captured.

These findings extend those of Russo et. al. (2018), who found that the largest components of neural activity during cycling (the top two principal components, which together captured ~35% of the neural variance) did not resemble velocity. Yet kinematic signals could still have been sizeable. As pointed out by Jackson (2018) and acknowledged in Russo et al. (2018), even the dominant signals could potentially reflect a joint representation of kinematic signals and their derivatives (e.g., vertical hand position and velocity). Furthermore, a signal can be sizeable even if it is not isolated in the top two principal components; indeed, the majority of variance lies outside those two dimensions. The present data reveal that both velocity- and position-correlating signals are very small – 10-fold smaller than the top two PCs. The fact that kinematic-correlating signals are small during some tasks (cycling) but sizeable in others (reaching) argues that they are most likely incidental.

Of course, even an incidental correlation could be useful. Yet the fact that kinematic-correlating neural dimensions are low-variance makes them a challenging substrate for decoding. For example, we identified a dimension where the projection of trial-averaged neural activity correlated with angular velocity, which is conveniently proportional to the quantity we wish to decode (velocity of virtual self-motion). However, because that dimension captured relatively little variance (1.2% ± 0.2% of the overall population variance; standard error across three sessions) the relevant signal was variable on single trials (**Fig. 2c**) and the correlation with angular velocity was poor. Generalization R^2^ was 0.63 for the segment of data shown, and was even lower when all data were considered (R^2^ = 0.51 ± 0.15, standard error across three sessions). Any decode based on this signal would result in many false starts and false stops. This example illustrates a general property: single-trial variability is expected for any low-variance signal estimated from a limited number of neurons.

We considered an alternative kinematic decoding strategy that more closely mimics prior successful reach-based approaches: decoding horizontal and vertical hand position. We chose position rather than velocity (which typically dominates reach-BMI decoding) because correlations with position were slightly stronger (compare *orange symbols* in **Fig. 2b**). A reasonable strategy would be to convert a two-dimensional position decode (**Fig. 2d**) into a one-dimensional self-motion command. Possible approaches for doing so include deciphering where on the circle the hand is most likely to be at each moment, computing the angular velocity of the neural state, or computing its angular momentum. However, consideration of such approaches raises a deeper question. If we are willing to non-linearly decode a one-dimensional quantity (self-motion) from a two-dimensional subspace, why stop at two dimensions? Furthermore, why not leverage dimensions that capture most of the variance in the population response, rather than dimensions that capture a small minority? Projections onto high-variance dimensions have proportionally less contribution from spiking variability. They are also less likely to be impacted by electrical interference or recording instabilities. While of little concern in a controlled laboratory environment, this is relevant to the clinical goals of prosthetic devices. One should thus, when possible, leverage multiple dimensions that jointly capture as much variance as possible.

We term this approach (**Fig. 2e**) task-specific subspace decoding, because it seeks one or more subspaces that capture robust response features as the subject attempts to move within the context of a particular task (in the present case, while generating a rhythmic movement to produce self-motion). Both the population-level response features, and the subspaces themselves, may be specific to that task. The necessity of task-specific decoding is already well-recognized when tasks are very different (e.g., reaching versus speaking) but may be beneficial even within the context of different tasks normally performed with the same limb.

To pursue this strategy, we identified three high-variance subspaces. The first was spanned by four ‘rotational’ dimensions (two each for forward and backward cycling) which captured elliptical trajectories present during steady-state cycling (Russo et al., 2018). The second was a single ‘moving-sensitive’ dimension, in which the neural state distinguished whether the monkey was stopped or moving regardless of movement direction (Kaufman et al., 2016). The third was a triplet of ‘initial-direction’ dimensions, in which cycling direction could be transiently distinguished in the moments after cycling began.

In subsequent sections we document the specific features present in these high-variance subspaces. Here we concentrate on the finding that the eight-dimensional space spanned by these dimensions captured 70.9% ± 2.3% of the firing-rate variance (**Fig. 2f**). Note that because dimensions are not orthogonal, the total captured variance is not the sum of that for each dimension. The ‘initial-direction’ dimensions, for example, overlap considerably with the ‘rotational’ dimensions. Nevertheless, the total captured variance was only modestly less than that captured by the top eight PCs (which capture the most variance possible), and much greater than that captured by spaces spanned by dimensions where activity correlated with kinematics and/or muscle activity (**Fig. 2b**). We thus based our BMI decode on activity in these eight high-variance dimensions.

The number of dimensions was chosen not to capture a particular proportion of neural variance, but was simply the natural result of focusing on high-variance neural features that possessed useful relationships with parameters a subject would wish to control in this task (e.g., cycling direction). Those neural features spanned eight dimensions that captured 70.9% of the variance. It is likely that, within the remaining 29.1% of the variance, there exist additional features that could be leveraged in other ways. We chose not to pursue that possibility as BMI performance (documented below) was already excellent. Still, decoding was not perfect and it is thus worth noting that further improvements may be possible.

### Direction of steady-state movement inferred from rotational structure

The dominant feature of the population neural response during steady-state cycling was a repeating elliptical trajectory (Russo et al., 2018). Thus, the core of our decoder was built upon this feature, leveraging the fact that forward-cycling and backward-cycling trajectories occurred in non-identical subspaces (**Fig. 3a**). We employed an optimization procedure to find a two-dimensional ‘forward plane’ that maximized the size of the forward trajectory relative to the backward trajectory. We similarly found an analogous ‘backward plane’. Planes cleanly captured rotations of the trial-averaged neural state (**Fig. 3a**) and, with filtering (*Methods*), continued to capture rotational features on individual trials (**Fig. 3b**). Forward and backward trajectories were not perfectly orthogonal. Nevertheless, the above procedure identified orthogonal planes where strongly elliptical trajectories were present for only one cycling direction. The data in Figure 3a,b are from a manual-control session (which employed multiple distances) to illustrate this design choice. In practice, planes were found based on decoder-training trials (7-cycle only) after trial-averaging.

**Figure 3.**
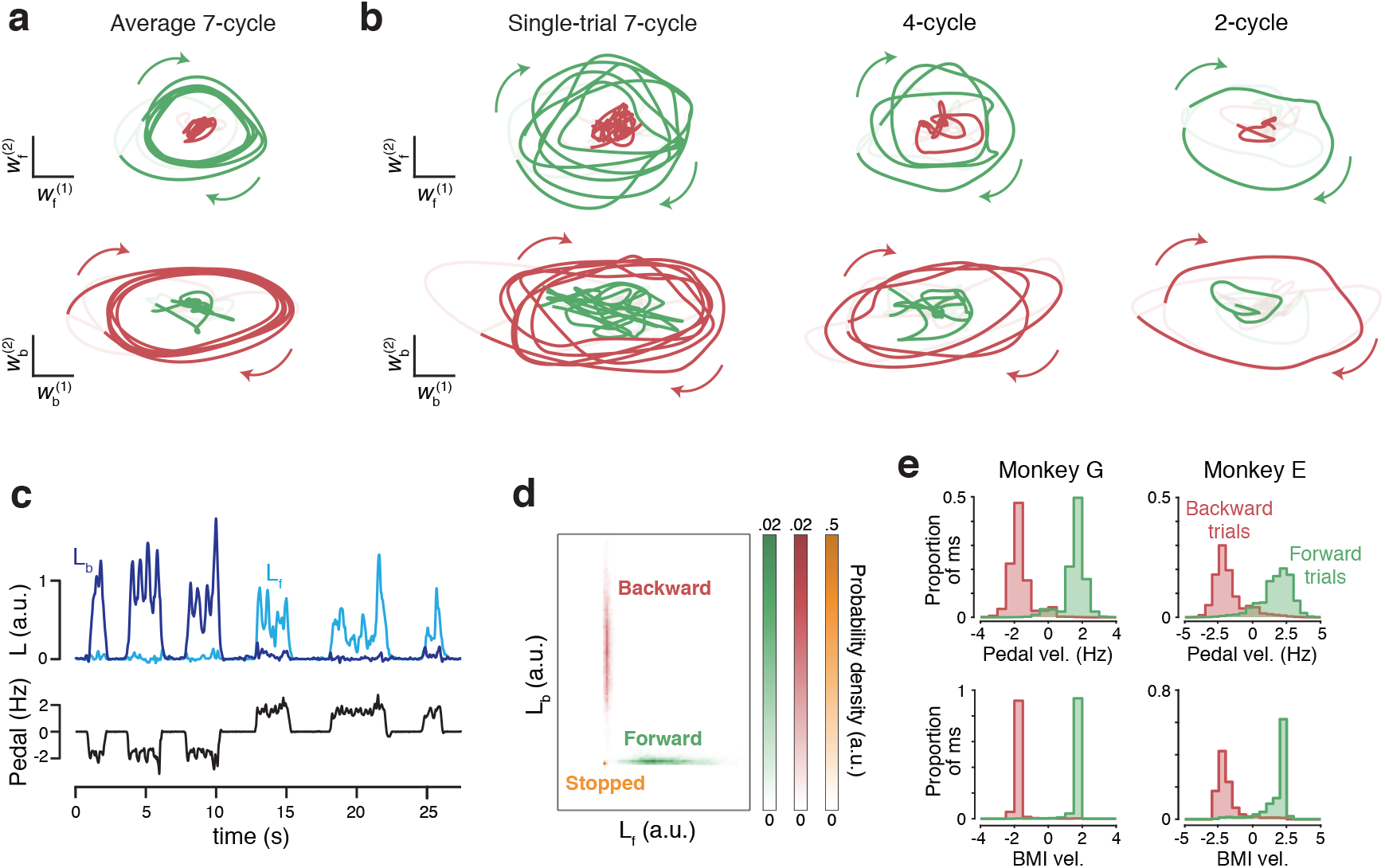
Leveraging rotational trajectories to decode velocity. **(a)** Trial-averaged population activity, during a manual-control session, projected onto the forward (*top*) and backward (*bottom*) rotational planes. Data are from seven-cycle forward (*green*) and backward (*red*) conditions. By design, the forward plane primarily captures rotational trajectories during forward cycling, and vice versa. Boldly colored portions of each trace highlight rotations during the middle cycles (a period that excludes the first and last half cycle of each movement). Colored arrows indicate rotation direction. Light portion of each trace corresponds to the rest of the trial. In addition to smoothing with a causal filter, neural data have been high-pass filtered to match what was used during BMI control. Data are from monkey G. **(b)** As in panel (a), but for six example single trials, one for each of the three distances in the two dimensions. **(c)** Example angular momentum (L) in the backward plane (*dark blue*) and forward plane (*bright blue*) during six trials of BMI control. Velocity of the pedal is shown in black. Although the pedal was disconnected from the task, this provides a useful indication of how the monkey was intending to move. Data are from the same day shown in panels **a** and **b**. **(d)** Probability densities of angular momentums found from the decoder training trials collected on the same day. **(e)** Histograms of BMI-control velocity (*bottom*) and (disconnected) pedal velocity (*top*) for all times the decoder was in the STEADY state (see Methods), across all BMI-control sessions.

To convert the four-dimensional neural state to a one-dimensional decode of self-motion, we compared angular momentum (the cross product of the state vector with its derivative) between the two planes. When moving backward (first three cycling bouts in **Fig. 3c**) angular momentum was sizeable in the backward plane (*dark blue*) but not the forward plane (*bright blue*). The opposite was true when moving forward (subsequent three bouts). Using the decoder-training trials, we considered the joint distribution of forward-plane and backward-plane angular momentum. We computed distributions when stopped (**Fig. 3d**, *orange*), when cycling forward (*green*) and when cycling backward (*red*) and fit a Gaussian to each. During BMI control, we computed the likelihood of the observed angular momentums under each of the three distributions. If likelihood under the stopped distribution was high, decoded velocity was zero. Otherwise, likelihoods under the forward and backward distributions were converted to a virtual velocity that was maximal when one likelihood was much higher and slower when likelihoods were more similar. Maximum decoded velocity was set based on the typical virtual velocity under manual control, when cycling at ~2 Hz.

Distributions of decoded velocity when moving under BMI control (**Fig. 3e**, *bottom*) were similar to the distributions of velocity that would have resulted were the pedal still operative (**Fig. 3e**, *top*). Importantly, distributions overlapped little; the direction of decoded motion was almost always correct. Decoded velocity was near maximal at most times, especially for monkey G. The decoded velocity obtained from these four dimensions constituted the core of our decoder. We document the performance of that decoder in the next section. Later sections describe how aspects of the decoder were fine-tuned by leveraging the remaining four dimensions.

### Performance

Monkeys performed the task very well under closed-loop BMI control (**Fig. 4** and **Movie 1**). Monkeys continued to cycle as normal, presumably not realizing that the pedal had been disconnected from the control system. The illusion that the pedal still controlled the task was supported by a high similarity between decoded virtual velocity and intended virtual velocity (i.e., what would have been produced by the pedal were it still controlling the task). The cross-correlation between these was 0.93 ± .02 and 0.81 ± .03 (monkey G and E, mean ± SD across sessions) with a short lag: 76 ± 4 ms and 102 ± 7 ms (**Fig. 4a**). There were also few false starts; it was exceedingly rare for decoded motion to be non-zero when the monkey was attempting to remain stationary on top of a target. False starts occurred on 0.29% and 0.09% of trials (monkeys G and E), yielding an average of 1.9 and 0.12 occurrences per day. This is notable because combatting unintended movement is a key challenge for BMI decoding. The above features – high correlation with intended movement, low latency, and few false starts – led to near-normal performance under BMI control (**Fig. 4b,c**). Success rates under BMI control (**Fig. 4d**, *magenta symbols*) were almost as high as under manual control (*open symbols*), and the time to move from target to target was only slightly greater under BMI control (**Fig. 4e**).

**Figure 4.**
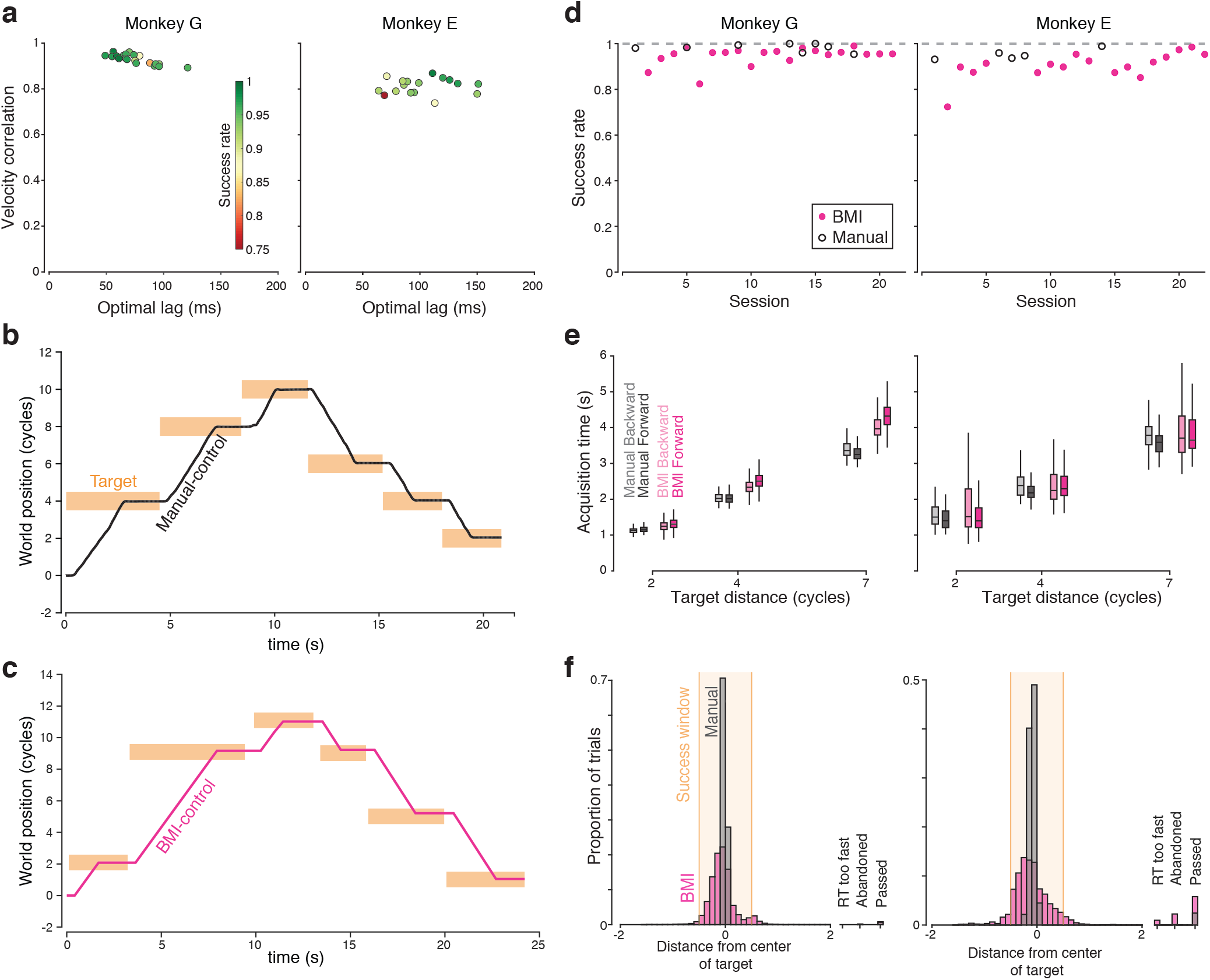
Decoder performance. **(a)** Summary of the cross-correlation between decoded virtual velocity under BMI control, and the virtual velocity that would have been produced by the pedal (which monkeys continued to manipulate normally). Each symbol corresponds to one BMI-control session, and plots the peak of the cross-correlation versus the lag where that peak occurred. Colors indicate success rate during that session. **(b)** Example manual-control performance for six consecutive trials, 3 forward and 3 backward. World position is expressed in terms of the number of cycles of the pedal needed to move that distance. For plotting purposes, the position at the beginning of this stretch of behavior was set to zero. Bars indicate the time that targets turned on and off (horizontal span) and the size of the acceptance window (vertical span). **(c)** Similar plot during BMI control. For ease of comparison, world position is still expressed in terms of the number of physical cycles that would be needed to travel that far, although physical cycling no longer had any impact on virtual velocity. **(d)** Success rate for both monkeys. Each symbol plots, for one session, the proportion of trials where the monkey successfully moved from the initial target to the final target, stopped within it, and remained stationary until reward delivery. Dashed line at 1 for reference. **(e)** Target acquisition times for successful trials. Center lines indicate median, the box edges indicate the first and third quartiles, and the whiskers include all non-outlier points (points less than 1.5 times the interquartile range from the box edges). Data are shown separately for the three target distances. **(f)** Histograms of stopping location from both monkeys. Analysis considers both successful and failed trials. The bars at far right indicate the proportion of trials where the monkey failed for reasons other than stopping accuracy per se. This included trials where monkeys disrespected the reaction time limits, abandoned the trial before approaching the target, or passed through the target without stopping.

Although the BMI decoder was trained using only data from 7-cycle movements, monkeys successfully used it to perform random sequences with different distances between targets (specifically, random combinations of 2, 4, and 7 cycles). For example, in Figure 4c the monkey successfully performs a sequence involving 2-, 7-, and 2-cycle forward movements, followed by 2-, 4-, and 4-cycle backward movements. The only respect in which BMI control suffered noticeably was accuracy in stopping on the middle of the target. Under manual control, monkeys stopped very close to the target center (**Fig. 4f,** *gray histogram*), which always corresponded to the ‘pedal-straight-down’ position. Stopping was less accurate under BMI control (*magenta histogram*). This was partly due to the fact that because virtual self-motion was swift, small errors in decoded stopping time became relevant. E.g., a 100 ms error corresponded to ~0.2 cycles of physical motion. The average standard deviation of decoded stopping time (relative to actual stopping time) was of this order: 133 (monkey G) and 99 ms (monkey E). The relatively larger error in BMI-control trials was also due to an incidental advantage of manual control: the target center was aligned with the pedal-straight-down position, a fact which monkeys leveraged to stop very accurately in that position. This strategy was not available during BMI control because the correct moment to stop rarely aligned perfectly with the pedal-straight-down position (this occurred only if decoded and intended virtual velocity matched perfectly when averaged across the cycling bout).

Performance was modestly better for monkey G versus E. This was likely due to the implantation of two arrays rather than one. Work ethic may also have been a factor; monkey E performed fewer trials under both BMI and manual control. Still, both monkeys could use the BMI successfully starting on the first day, with success rates of 0.87 and 0.74 (monkey G and E). Monkey G’s performance rapidly approached his manual-control success rate within a few sessions. Monkey E’s performance also improved quickly, although his manual-control and BMI-control success rates were mostly lower than Monkey G’s. The last five sessions involved BMI success rates of 0.97 and 0.96 for the two monkeys. This compares favorably with the overall averages of 0.98 and 0.95 under manual control. Although this performance improvement with time may relate to adaptation, the more likely explanation is simply that monkeys learned to not be annoyed or discouraged by the small differences in decoded and intended velocity.

### State machine

The performance documented above was achieved using a state-dependent decode (**Fig. 5**). The rotational-feature-based strategy (described above) determined virtual self-motion in the “STEADY” state. Thus, self-motion could be forward, backward, or stopped while in STEADY. The use of other states was not strictly necessary but helped fine-tune performance. State transitions were governed by activity in the moving-sensitive dimension, which was translated into a probability of moving, *p_move_*, computed as described in the next section. If *p_move_* was low, the STOP state was active and decoded virtual velocity was enforced to be zero. For nearly all times when the STOP state was active, the angular-momentum-based strategy would have estimated zero velocity even if it were not enforced. Nevertheless, the use of an explicit STOP state helped nearly eliminate false starts.

**Figure 5.**
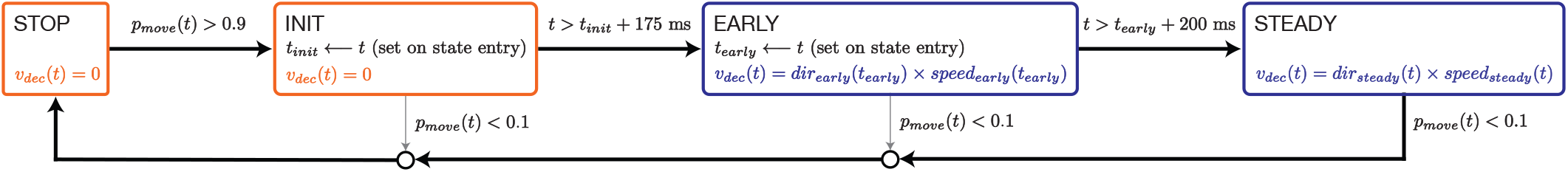
State machine diagram. BMI motion was determined by a state machine with four states: STOP, INIT, EARLY, and STEADY, corresponding to the different stages of a typical trial. The output of the state machine at every millisecond was an estimate of decoded velocity through the virtual environment, *v_dec_*, which was then smoothed and integrated to compute virtual position. Black arrows indicate the typical path of a successful BMI trial and gray arrows indicate all other possible transitions. State transitions were governed by activity in the moving-sensitive dimension, which was translated into a probability of moving, *p_move_*. While *p_move_* was low, the STOP state was active and decoded velocity was set to zero. When *p_move_* became high, the INIT state was entered but decoded velocity remained zero. If *p_move_* remained high for 175 ms, the EARLY state was entered and velocity was decoded using the initial-direction dimensions. After another 200 ms, the STEADY state was entered and decoded velocity depended on the neural state in the rotational dimensions. If *p_move_* dropped below 0.1 at any point, STOP was reentered. States in which progress is made through the virtual environment are highlighted in blue and states in which BMI motion is held at zero are highlighted in orange.

The use of a STOP state was also useful because exiting STOP indicated a likely transition to moving, allowing us to leverage the initial-direction dimensions. When *p_move_* became high, the INIT state was entered but decoded velocity remained zero. After 175 ms, the EARLY state was entered and velocity was decoded using the initial-direction dimensions (see below). After an additional 200 ms, the STEADY state was entered. The decode then depended only on the four rotational dimensions. Values of *p_move_* < 0.1 always produced a transition back to STOP.

This typically occurred from STEADY to STOP as the movement successfully terminated. However, it could also occur from the other two states. This was especially helpful if, when stopped, *p_move_* became high only briefly (and thus presumably erroneously). In such cases the state transitioned from INIT back to STOP with decoded velocity never departing from zero.

### Inferring the probability of moving

Decoders that directly translate neural state to cursor velocity have historically experienced difficulty remaining stationary when there is no intended movement. The ability to successfully decode stationarity is of even greater importance for self-motion. This was central to our motivation for using a state machine with distinct stopped and moving states (Ethier et al., 2011; Kao et al., 2017a; Kemere et al., 2008). To govern state transitions, we leveraged the moving-sensitive dimension: the dimension where neural activity best discriminated whether the monkey was moving versus stopped, identified using linear discriminant analysis. Projecting trial-averaged data onto that dimension (**Fig. 6a**) revealed a transition from low to high just before movement onset, and back to low around the time movement ended. This pattern was remarkably similar regardless of cycling direction (*red* and *green* traces largely overlap). Activity in this dimension behaved similarly for single trials (**Fig. 6b**). The data in Figure 6a,b are from a manual-control session, and illustrate why activity in this dimension was considered useful. Before BMI decoding, the moving-sensitive dimension was identified based on the 50 decoder-training trials, and we considered the distribution of activity in that dimension when stopped (**Fig. 6c**, *orange*) and moving (*blue*). We fit a Gaussian distribution to each, which was then used during BMI-control when computing the probability of moving, *p_move_*.

**Figure 6.**
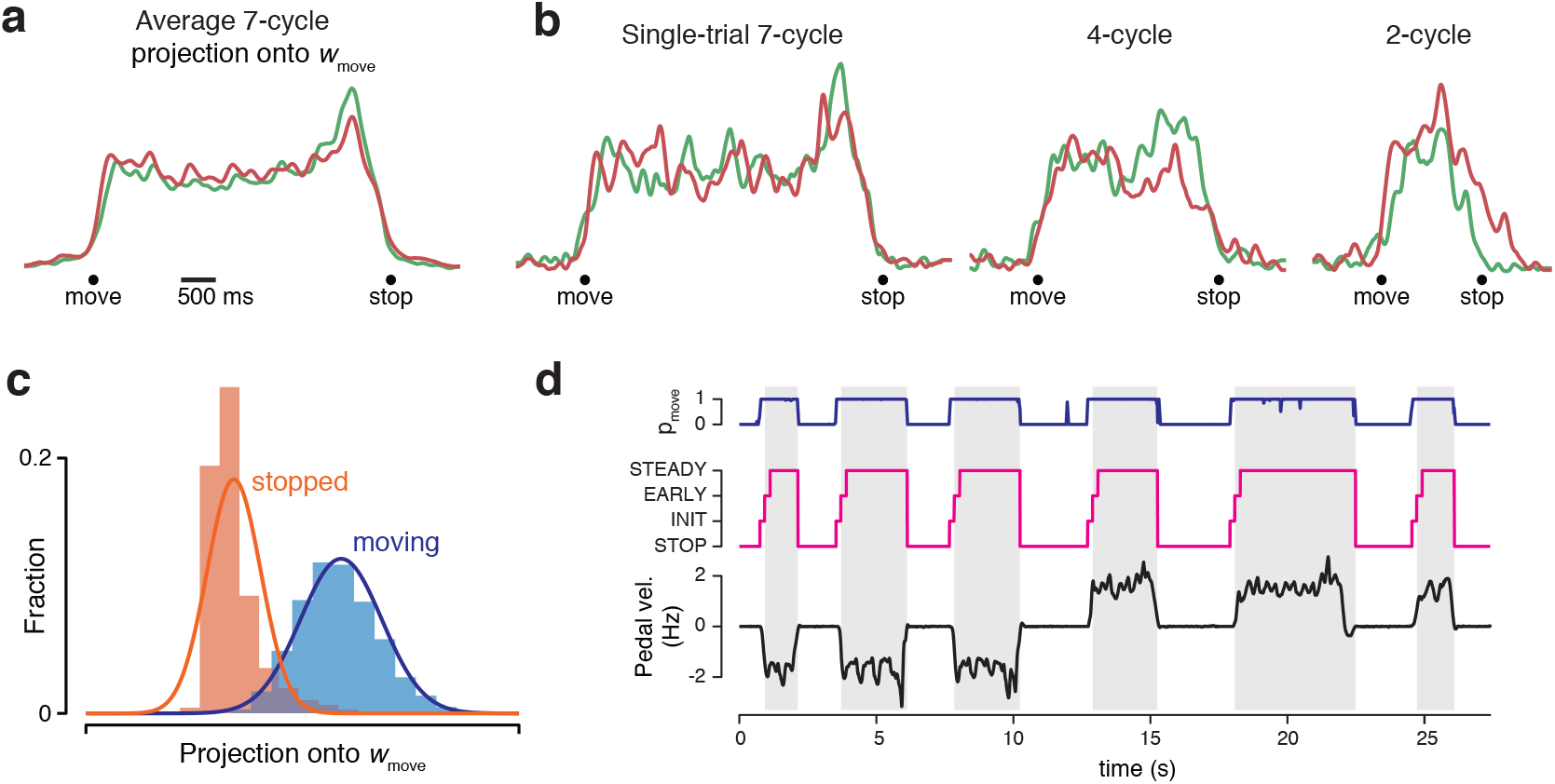
Leveraging the moving-sensitive dimension to infer probability of moving. **(a)** Trial-averaged population activity, during a manual-control session, projected onto the moving-sensitive dimension (same session and trials as Figure 3a). **(b)** As in panel (a), but for six example single trials (same trials as in Figure 3b). **(c)** Histogram of the neural state projected onto the moving-sensitive dimension for decoder training data. The neural state was measured every ten milliseconds, at times when the monkey was stopped within a target (*orange*) or actively cycling (*blue*). Traces show Gaussian fits used to compute *p_move_*. **(d)** Example time-course, during BMI control, of *p_move_* (*blue*) and the active state (*magenta*). Gray regions show times when the decoder produced virtual movement (i.e., when in EARLY or STEADY). These times corresponded well to times when the monkey was intending to move, as indicated by the angular velocity of the disconnected pedal (*black*). Note also that transient inappropriate spikes in *p_move_* (as seen here around 12 s) do not lead to false starts because either they don’t exceed 0.9, as was the case here, or they are too brief and the EARLY state is never reached. Same example data as in Figure 3c.

To compute *p_move_*, we used a Hidden Markov Model (HMM) (Kao et al., 2017a; Kemere et al., 2008) allowing the current estimate of *p_move_* to depend on all prior observations. Doing so allowed the computation of *p_move_* to be robust with respect to the modest overlap in the stopped and moving distributions. An HMM can ignore transient weak evidence for moving while still transitioning swiftly given strong evidence. As a technical aside, because the HMM assumes observations are independent, we did not use filtered rates (which were used for all other aspects of the decode) but instead considered spike counts in non-overlapping 10 ms bins. This was true both when determining the moving-sensitive dimension and while under BMI-control.

During BMI control, *p_move_* (**Fig. 6d**, *blue*) was typically near unity during intended movement (i.e., when the monkey was actually cycling, *black*) and near zero otherwise. The value of *p_move_* determined transitions both out of STOP and into STOP (**Fig. 5**). We employed a conservative design; entering EARLY (the first state that produced virtual movement) required that *p_move_* exceed 0.9 and remain consistently above 0.1 for 175 ms. This led to a very low rate of false starts (~2 per day for monkey G and ~1 every ten days for monkey E). The transition to EARLY (**Fig. 6d**, left edge of *gray regions*) occurred on average 117 and 194 ms after physical movement onset (monkeys G and E). Trial-to-trial variability around these mean values was modest: standard deviations were 93 and 138 ms (computed within session and averaged across sessions). As discussed above, estimated stopping time (when *p_move_* dropped below 0.1) was also decoded with only modest trial-to-trial variability.

### Inferring initial movement direction

Angular momentum of the neural state in the forward and backward planes became substantial a few hundred milliseconds after *p_move_* became high. Thus, the EARLY state became active before the direction of movement could be inferred from the elliptical trajectories. To overcome this problem, we leveraged the three-dimensional initial-direction subspace. The initial-direction subspace comprised the top three PCs found from portions of the training data surrounding movement onset (a 200 ms segment from each trial, beginning at the time the decoder would enter the INIT state, see *Methods*). The difference between the trajectory on forward and backward trials began to grow just prior to physical movement onset, both on average (**Fig. 7a**) and on individual trials (**Fig. 7b**; *solid* trajectory segment shows −200 to +175 ms relative to movement onset). For each of the 50 decoder-training trials, we considered the neural state in these dimensions, measured 175 ms after decoded movement onset (**Fig. 7c**). We fit Gaussian distributions separately for forward and backward trials.

**Figure 7.**
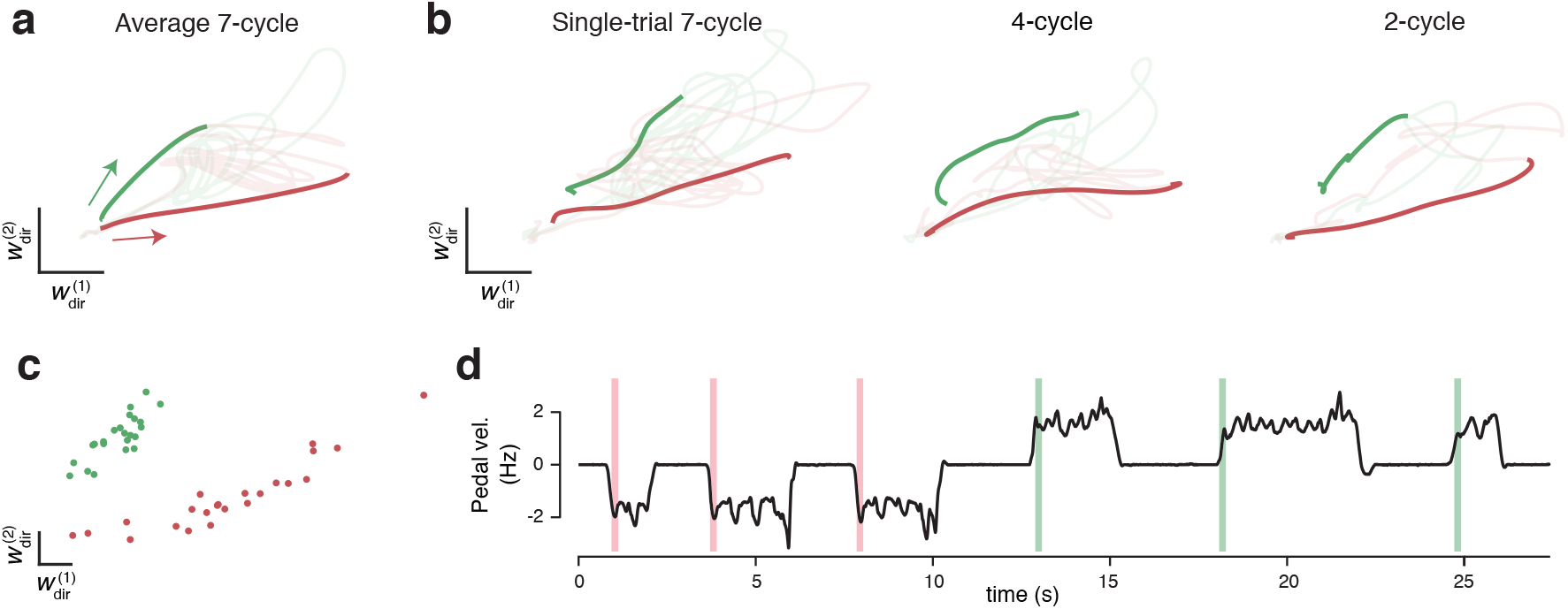
Leveraging initial-direction dimensions to allow low-latency decoding. **(a)** Trial-averaged population activity, during a manual-control session, projected onto two (of three) initial-direction dimensions (same session and trials as Figure 3a and 6a). Boldly colored portions of traces highlight −200 ms to +175 ms relative to physical move onset. Arrows indicate direction of trajectories. **(b)** As in panel (a), but for six example single trials (same trials as in Figure 3b and 6b). **(c)** The location of the neural state, for decoder training data, at the time the state-machine (applied post-hoc to that training data) entered the EARLY state. This data (50 total trials) was used to fit two Gaussian distributions. During BMI control, when the EARLY state was entered, virtual direction was determined by which distribution maximized the data likelihood. **(d)** Example of initial-direction decoding during BMI control. Colored windows show the times in the EARLY state, with red and green indicating decoded direction. Same example data as in Figure 3c and 6d.

During BMI control, upon transition from INIT to EARLY, we computed the likelihood of the neural state under each distribution. A simple winner-take-all computation determined the direction of virtual velocity during the EARLY state. The inference of movement direction during EARLY was correct on 94% and 82% of trials (monkeys G and E). After 200 ms, the STEADY state was entered and virtual velocity was controlled thereafter by activity in the rotational dimensions. **Figure 7d** illustrates moments (*colored regions*) where the EARLY state was active and determined virtual motion (physical pedal velocity is shown for reference). These moments were brief, and thus produced only a very modest improvement in time to reach the target. However, we still employed this strategy because our goal was to build a BMI decode that closely tracked intended movement and felt responsive to the subject.

### Speed control

The excellent performance of the decoder was aided by the relative simplicity of behavior: when monkeys moved, they did so at a stereotyped speed. This allowed us to concentrate on building a decode algorithm that decoded intended direction with accurate timing, and remained stationary if movement was not intended. However, that decode provided only limited control of movement speed. An obvious extension is to allow finer-grained speed control. This would presumably be desired by users of a self-motion BMI. Furthermore, speed control provides one possible way of steering: e.g., by decoding the relative intensity of intended movement on the two sides of the body. While we do not attempt that here, we still considered it important to determine whether the neural features we identified could support speed control.

That assessment required a task where speed control is necessary for success. We thus trained one monkey to track various speed profiles as he progressed through the virtual environment. Two floating targets were rendered in the foreground as the monkey cycled. The distance between them reflected the difference between actual and instructed speed. Obtaining juice required aligning the two floating targets while progressing towards a final target, on which he stopped to obtain additional reward. The task was divided into trials, each of which required moving a distance equivalent to twenty cycles under manual control. We used eight trial-types, four each for forward and backward cycling. Two of these employed a constant target speed (equivalent to 1 or 2 Hz cycling) and two involved a ramping speed (from 1 Hz to 2 Hz or vice versa) (**Fig. 8a,b**). The decoder was trained based on a small number of decoder-training trials, employing only the two constant speeds, performed at the beginning of each BMI-control session. Performance comparisons were made with manual-control sessions that employed all conditions.

**Figure 8.**
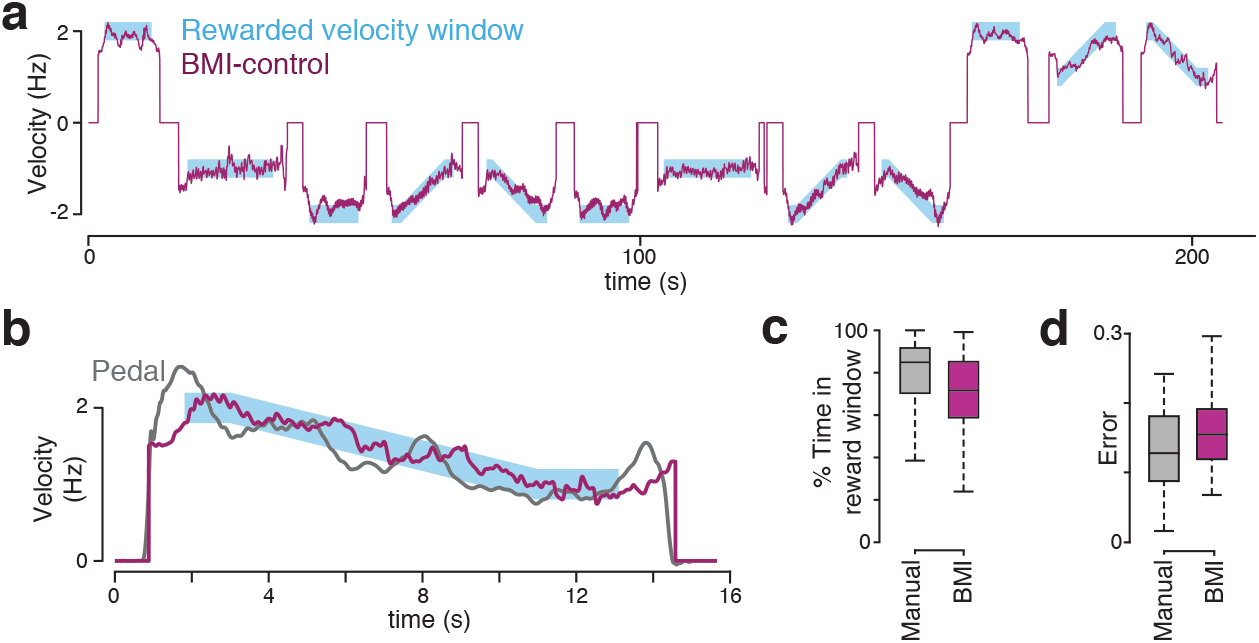
Performance in the modified task requiring speed tracking. **(a)** Instructed velocity and BMI-decoded virtual velocity during 12 contiguous trials of BMI control. **(b)** Expanded view of one example trial (the last trial from panel a). The virtual velocity that would have been produced by the pedal is shown in gray for comparison. **(c)** Percentage of time spent in rewarded velocity window for trials in manual-control (2 sessions, 333 trials) and BMI-control (3 sessions, 349 trials). Center lines indicate median, the box edges indicate the first and third quartiles, and the whiskers include all non-outlier points (points less than 1.5 times the interquartile range from the box edges). **(d)** Mean absolute error (MAE) between instructed velocity and virtual velocity for both manual control and BMI control sessions. One mean error was computed per trial. Same format as (c).

Our decode strategy was largely preserved from that described above. However, we used a modified state machine (**Fig. 9**) and a slightly different algorithm for transforming rotations of the neural state into decoded virtual velocity. Direction was still determined based on which distribution (forward or backward) produced the higher likelihood of observing the measured angular momentums (as in **Fig. 3d**). Once that choice was made, speed was determined by the angular velocity of the neural state in that plane. Thus, faster rotational trajectories led to faster decoded virtual velocity. We chose a scaling factor so that a given neural angular velocity produced the speed that would have been produced by physical cycling at that angular velocity. Neural angular velocity was exponentially filtered with a time constant of 500 ms. The filter memory was erased on entry into a movement state (EARLY or STEADY) from a stopped state (INIT or EXIT) to allow brisk movement onset (see *Methods)*.

**Figure 9.**
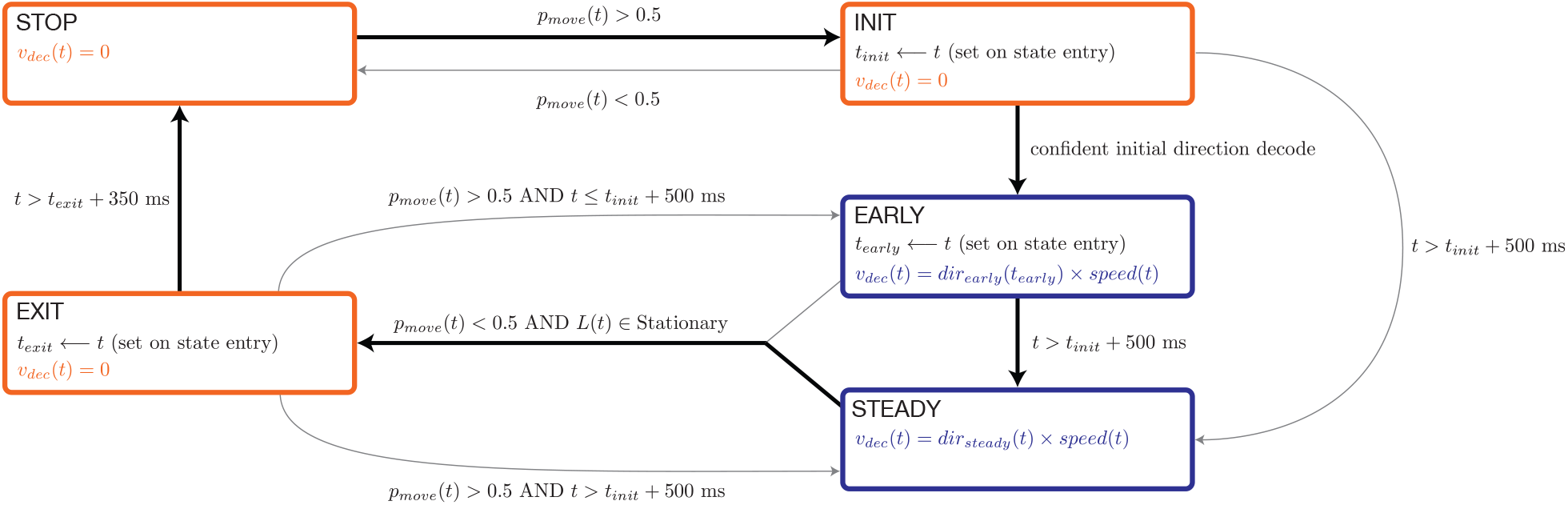
State machine diagram for speed-control experiments. BMI motion for the speed-tracking experiment was determined by a state machine with five states: STOP, INIT, EARLY, STEADY, and EXIT. Each of these states describes how to compute virtual velocity, *v_dec_*, which gets integrated every millisecond into virtual position. Black arrows indicate the typical path of a successful BMI trial (gray arrows indicate all other transitions) and colors differentiate states in which there is progress through the virtual environment (blue) from states in which BMI motion is zero (orange). In the STOP, INIT, and EXIT states, virtual velocity is simply set to zero. During the EARLY and STEADY states, virtual velocity is computed as the product of decoded direction and decoded speed, which is computed based on activity in the rotational dimensions. Transitions between the states were determined by *p_move_* (computed from the moving-sensitive dimension), angular momentum vector L (computed from the rotational planes), and the neural state in the initial-direction dimensions. Details on computing each of the neural features used in this state machine are provided in the section ‘Neural features for speed-tracking’. While in the STOP state, virtual velocity remains zero. If *p_move_* goes high, the INIT state is entered (time of entry is denoted *t_init_*), but virtual velocity remains zero. From the INIT state, one of three things can happen: 1) *p_move_* goes low and the movement aborts with a transition back to the STOP state, 2) the neural state in the initial-direction dimensions becomes sufficiently close to the learned distributions of neural states for forward or backward conditions, triggering a transition to the EARLY state, or 3) no transition occurs within 500 ms of *t_init_*, triggering a transition to the STEADY state. The INIT state serves to withhold BMI motion in situations where there is a transient spike in *p_move_* and to delay movement when the early estimate of decoded direction is only weakly confident. The EARLY state dictates how virtual velocity is computed in the early portion of movement (when less than 500 ms have elapsed since *t_init_*). The STEADY state determines virtual velocity for the remainder of decoded movement. While in the EARLY or STEADY states, if *p_move_* goes low and the angular momentums in the rotational planes are consistent with a stationary pedal, a transition will occur to the EXIT state and virtual velocity will again go to zero. After 350 consecutive ms in the EXIT state, the state machine transitions back to the STOP state. However, if *p_move_* goes high while in the EXIT state, a transition back to either EARLY or STEADY will occur. The transitions from EXIT back to EARLY and STEADY exist to allow a rapid return to decoded movement if neural activity briefly, erroneously reflects that the monkey is ceasing movement, but then returns to exhibiting robust activity consistent with an ongoing movement.

The above strategy allowed smooth BMI control of movement speed. In fact, it tended to give BMI control an intrinsic advantage over manual control. In manual control, the angular velocity of the pedal was naturally modulated within each cycle (being higher on the downstroke), resulting in a fluctuating virtual velocity. Such fluctuations mildly impaired the ability to match target speed under manual control. To allow a fair comparison, we thus also applied an exponential filter to virtual velocity under manual control. Filters were chosen separately for BMI (τ = 500 ms) and manual control (τ = 1000 ms) to maximize performance. This was done informally, in the earliest session, by lengthening the filter until success rate roughly plateaued. The filter then remained fixed for all further sessions.

Under BMI control, decoded virtual speed closely tracked instructed speed (**Fig. 8a,b** and **Movie 2**). This was true both for constant speeds and when speed modulated with time, even though the decoder was not trained on such modulations. To compare BMI with manual control (which were performed on separate days) we considered all trials where the monkey completed the portion of the trial that required matching speed (87% of trials in arm control, and 79% in BMI control). The monkey was able to match instructed speed nearly as accurately under BMI control as under manual control. This was true judged both by time within the rewarded speed window (**Fig. 8c**) and by the error between virtual and instructed velocity (**Fig. 8d**).

## Discussion

### Task-specific subspaces

The lack of high-variance kinematic-correlating dimensions is consistent with Russo et al. (2018) which found that neural trajectories in the top two dimensions were inconsistent with a representation of muscle activity or hand velocity. Yet the findings of Russo et al. did not rule out the possibility that kinematic-correlating dimensions might still be reasonably high-variance even if they were not dominant. This is true during reaching, where the top two dimensions do not correlate well with kinematics (Kaufman et al., 2016) but other dimensions do. Furthermore, a combined representation of velocity and position was a plausible, if unappealing, explanation for the dominant circular trajectories (Jackson, 2018). In fact, dimensions where activity correlated well with kinematics contained very little variance: ~1-4% per dimension. The same was true of dimensions that correlated with muscle activity.

The lack of dimensions where activity consistently correlates with external variables accords with the hypothesis that the largest signals in motor cortex are not ‘representational’ – they do not encode variables but are instead essential for noise-robust dynamics (Churchland et al., 2012; Russo et al., 2018; Seely et al., 2016; Sussillo et al., 2015). Those dynamics produce outgoing commands that *are* representational (they covary with the variables they control) but are low-variance. Because the dominant neural signals must be produced by noise-robust dynamics, they obey the property of low trajectory tangling: similar neural states avoid being associated with dissimilar derivatives (Russo et al., 2018). There is little guarantee that the way in which low tangling is achieved will be the same across tasks. On the contrary, the task often determines the most natural way to keep tangling low. Thus, the identity and number of high-variance dimensions will likely vary across tasks, as will the basic response features captured by those dimensions. This possibility is supported by network models that perform multiple tasks (Duncker et al., 2020; Flesch et al., 2021; Logiaco et al., 2019) or subtasks (Zimnik and Churchland, 2021). When different tasks require very different dynamics, a very natural way to ‘switch’ dynamics is to alter the occupied subspace. If this is indeed a common strategy in the nervous system, both the intrinsic covariance structure of the neural activity, and the nature of its correlations with kinematics, will likely be task specific.

While task specificity presents practical challenges for decoders, it also affords opportunities and certainly does not preclude high-performance decoding. We identified subspaces very differently from most reach-based BMI approaches, and converted neural activity to movement using different (and often more non-linear) approaches. Yet we achieved BMI control that was sufficiently natural that monkeys appeared not to notice that the task was no longer under manual control. By most measures (success rate, time to target) performance under BMI control was remarkably close to that under manual control. The main limitation of BMI control was stopping accuracy. Although our algorithm detected stopping with ~ 0.1 second precision, even small discrepancies could lead to the target being over or undershot by a noticeable amount. A beneficial feature of our BMI decode is that it almost never produced movement when it was not intended. With rare exceptions, truly zero velocity was decoded when the monkey was intending to remain stopped on the target. We consider this a particularly important attribute of any self-motion decoding algorithm, due to the potentially large consequences of unintended movement of the whole body.

Perhaps the largest apparent drawback of task-specific decoders is that they do not offer the promise of across-task generalization, which has been a longstanding goal in BMI development. Our approach, being based on reliable but non-representational signals, remains subject to this drawback (no more or less so than traditional reaching decoders). However, within-task generalization is still very much possible. In the current work, training was on 7-cycle trials only and the decoder generalized to other trial lengths (arbitrary distances) since the rotational structure we leveraged mapped easily onto portions of cycles. Therefore, it was not the case that the decoder learned specifically to start, move a specified distance, and then stop. It was able to capture, on a millisecond-by-millisecond basis, the velocity of intended movement. Similarly, in the speed-tracking task variant, we trained only on two constant speed profiles (1 and 2 cycles/s) and were able to decode all observed speeds, demonstrating that the decoder could interpolate speed. Similar results could be achieved in other task situations, by leveraging the particular geometry of the dynamics; indeed, this is the mechanism by which center-out reaching decoders interpolate to produce untrained reach angles.

To develop a system that can effectively perform multiple tasks (e.g. self-motion, reaching, handwriting) with this approach, a new task-specific subspace decoder will be necessary for each task. While we cannot offer a concrete recipe for identifying the neural features that would best subserve decoding in a given task, being opportunistic in leveraging high-variance signals is likely to lead to success. Multiple decoder modules could then be combined. How this should be accomplished depends upon something not yet known: are the high-variance subspaces the same or different across tasks? If subspaces are the same (or overlap) that would complicate decoding; the decoder would have to decipher the task before knowing which features to leverage. If subspaces are fully orthogonal, then the features relevant to one decoder would conveniently fall in the null space of decoders for other tasks. A reasonable speculation is that subspaces will overlap when tasks are similar enough that correlations with to-be-decoded features are stable (Gallego et al., 2020), and will be nearly orthogonal when tasks are different enough that they require very different decoding approaches. Our work provides a hint that this may be true. The dimensions occupied during forward versus backward cycling were close enough to orthogonal that we could decode them independently and without confusion. Yet whether this is generally true remains to be seen. One could of course attempt to completely avoid such issues by focusing on the one subspace that must be preserved across tasks: that conveying the outgoing motor commands. However, as noted above this subspace may be very low variance, making it both difficult to identify and difficult to leverage without very large-scale simultaneous recordings.

### Population dynamics in decoding

A number of studies have modeled neural dynamics to improve online (Kao et al., 2015) or offline (Aghagolzadeh and Truccolo, 2016; Gallego et al., 2020; Kao et al., 2017b; Liu et al., 2019) decoding of movement kinematics. Although these studies assumed that their low-dimensional dynamics contained kinematic coding signals, they allowed for signals that did not correlate directly with kinematics to help infer those that do. In linear decoding, the value of a given variable depends upon the neural state in just one dimension: the dimension defined by the regression weights. Nevertheless, inferring the neural state in that dimension may benefit from a dynamical model that spans multiple dimensions. Much like the present approach, this allows the decoder to leverage features that are robust, even if they do not directly correlate with the kinematic parameters of interest. The present approach extends this idea to situations where there may be no high-variance dimensions that can be linearly decoded, and/or where the most prominent features are not well-described by linear dynamics. Accordingly, any method that allows us to better estimate the neural state stands to improve our results. Here, we used a linear Kalman filter, but other techniques (linear or nonlinear) could be considered in the future (e.g. LFADS (Pandarinath et al., 2018), Bayesian methods). Many such methods are presently acausal (e.g., LFADS employs knowledge of the future) but could adapted for use in real-time applications.

Another way of leveraging additional dimensions – or even the full neural state – would be to use a trained RNN to decode parameters of interest (Anumanchipalli et al., 2019; Sussillo et al., 2012; Willett et al., 2020). This is a promising approach, and avoids the need to hand-select features. However, it does not itself solve the problem of generalization. Through training, these networks can identify useful and noise-robust features, but can only do so for the patterns of neural activity included in the training datasets (e.g. a network trained to decode reaches would not be able to leverage the covariance structure during cycling).

### Self-motion BMIs

In choosing an alternative to traditional reach-based BMI tasks, we had three reasons for focusing on a BMI for virtual self-motion. First, our recently developed cycling task naturally lends itself to this application. Second, BMI-controlled self-motion is likely to be desired by a large patient population (potentially much larger than the population that desires BMI-controlled cursors or robot arms). Third, prior work has demonstrated that BMI control of self-motion is viable (Libedinsky et al., 2016; Rajangam et al., 2016). In particular, Rajangam et al. (Rajangam et al., 2016) demonstrated BMI control of a physical wheelchair based on neural activity recorded from monkey motor and somatosensory cortex. Our work supports their conclusion that BMI-controlled self-motion is possible, and demonstrates the feasibility of an alternative decode strategy. Rajangam et al. employed a traditional decode strategy: linear filters transformed neural activity into the key variables: translational and angular velocity. That strategy allowed monkeys to navigate ~2 meters to a target (which had to be approached with an accuracy of +/− ~0.2 meters, or 10% of the distance traveled) in an average of 27-49 seconds (depending on the monkey and degree of practice). In our task, monkeys had to stop with similar relative accuracy: +/− 0.5 cycles, or 7% of the distance traveled for a seven-cycle movement. They traversed those seven cycles in ~4 seconds under BMI control (averages of 4.3 and 3.7 seconds for monkey G and E). While this is roughly tenfold faster, we stress that movement durations are not directly comparable between our task and theirs. Success requirements differed in multiple ways. For example, Rajangam et al. required that monkeys turn *en route* (which adds considerable challenge) but did not require them to stop on the target location. Yet while direct comparison is not possible, a tenfold improvement in time-to-target argues that task-specific subspace decode strategies can be effective and should be explored further.

An obvious limitation of the current study is that we did not explore strategies for steering, which would be essential to a real-world self-motion prosthetic. There exist multiple candidate strategies for enabling steering. Rajangam et al. used a Wiener filter to decode angular velocity of the body. While straightforward, this strategy appears to have had limited success: even during training, the R^2^ of their angular velocity decode was 0.16 and 0.12 for the two monkeys. One alternative strategy would be to apply our decode strategy bilaterally, and employ a comparison (e.g., between left and right cycling speed) to control angular velocity. This strategy should be viable even with a unilateral implant; recent work has shown that information about both forelimbs can be decoded equally well from either hemisphere (Ames and Churchland, 2019; Heming et al., 2019). Another strategy would be to control translational velocity using the strategies developed here, but use a reach-like decode for steering (rather like pedaling a bicycle while also steering). Which (if any) of these three strategies is preferable remains a question for future experiments.

## Acknowledgements

We thank Y. Pavlova for expert animal care, A. Russo for sharing code and data for preliminary analysis, and E. Oby for surgical expertise and assistance. This work was supported by NINDS 1K99NS115919, NINDS 1DP2NS083037, NIH CRCNS R01NS100066, NINDS 1U19NS104649, the Simons Foundation (SCGB#325233 and SCGB#542957), the Grossman Center for the Statistics of Mind, the McKnight Foundation, P30 EY019007, a Klingenstein-Simons Fellowship, and the Searle Scholars Program.

## Notes

### Competing Interest Statement

The authors have declared no competing interest.

### Summary of Updates

Updated Abstract, Introduction, Results, Figure 2, and Discussion sections.

